# Role of Era in Assembly and Homeostasis of the Ribosomal Small Subunit

**DOI:** 10.1101/525360

**Authors:** Aida Razi, Joseph H. Davis, Yumeng Hao, Dushyant Jahagirdar, Brett Thurlow, Kaustuv Basu, Nikhil Jain, Josue Gomez-Blanco, Robert A. Britton, Javier Vargas, Alba Guarné, Sarah A. Woodson, James R. Williamson, Joaquin Ortega

## Abstract

To reveal the role of the essential protein Era in the assembly of the 30S ribosomal subunit, we analyzed assembly intermediates that accumulated in Era-depleted *Escherichia coli* cells using quantitative mass spectrometry, cryo-electron microscopy and in-cell footprinting. Our combined approach allowed for visualization of the small subunit as it assembled and revealed that with the exception of key helices in the platform domain, all other 16S rRNA domains were able to fold even in the absence of Era. Notably, the maturing particles did not stall while waiting for the platform domain to mature and instead re-routed their folding pathway to enable concerted maturation of other structural motifs spanning multiple rRNA domains. We also found that binding of Era to the mature 30S subunit destabilized helix 44 and the decoding center preventing binding of YjeQ, another assembly factor. This work establishes Era’s role in ribosome assembly and suggests new roles in maintaining ribosome homeostasis.

## INTRODUCTION

The bacterial 70S ribosome is made of the small (30S) and large (50S) subunits and consists of over fifty different ribosomal proteins (r-proteins) and ribosomal RNA (rRNA) that must fold and associate. During this assembly, the 16S rRNA in the 30S subunit and the 23S and 5S rRNA molecules in the 50S subunit fold according to energy landscapes comprised of multiple parallel assembly pathways (Adilakshmi et al., 2005; Talkington et al., 2005). *In vivo*, ribosome biogenesis is aided by subunit-specific protein assembly factors that enable rapid (~two minutes) and efficient ribosome biogenesis (Shajani et al., 2011). Although the specific function of most of these assembly factors is still unclear, they are generally thought to act by binding to the maturing particles and modifying their energy landscapes to favor folding pathways and to prevent the rRNA from falling into local energy minima (Talkington et al., 2005).

The ribosome assembly factor Era (*Escherichia coli* Ras-like) is known to participate in 30S subunit maturation, however its precise role is still largely unknown. This protein is universally conserved in both eukaryotes and prokaryotes (Leipe et al., 2002) and it is essential for both Gram-negative (Anderson et al., 1996; March et al., 1988; Takiff et al., 1989) and Gram-positive bacteria (Minkovsky et al., 2002; Sato et al., 1998). Era is comprised of a N-terminal GTPase domain and a C-terminal KH (K-homologue) domain connected by a 17 amino acid long flexible linker whose length is important for its function (Inoue et al., 2003). The GTPase domain consists of a central β-sheet flanked by five helices. The KH domain has a high structural similarity to the RbfA assembly factor and folds following a type 2 (αββααβ) KH folding pattern. This KH domain is necessary for Era to bind the 16S rRNA and the 30S subunit (Hang and Zhao, 2003; Johnstone et al., 1999).

Crystallography studies with purified Era and RNA fragments derived from the 3’ end of the 16S rRNA (Tu et al., 2009) revealed that the two domains of Era can adopt a ‘closed’ and an ‘open’ conformation. In the apo or GDP-bound states, Era adopts the open conformation in which the nucleotide binding site is accessible but the RNA binding site in the KH domain is occluded. Binding of GTP is thought to drive Era to the closed state, thus allowing for rRNA binding (Chen et al., 1999; Ji, 2016; Tu et al., 2009). According to this model, subsequent rRNA binding, which is known to stimulate GTP hydrolysis (Tu et al., 2009), would then revert Era to the open state and trigger release of Era from the rRNA. These findings are consistent with equilibrium binding assays in which Era exhibited increased affinity for rRNA in the presence of GDPNP, a non-hydrolyzable GTP mimic, relative to that in the presence of GDP (Sayed et al., 1999; Thurlow et al., 2016).

Although binding to the isolated rRNA fragment can be modeled by this two-state conformational switch model, Era may adopt additional conformations when bound to entire 30S subunit. Indeed, a low resolution cryo-electron microscopy (cryo-EM) structure of Era in complex with the 30S subunit found that neither of the aforementioned conformations were compatible with the orientation of the two domains of Era when the factor was bound to the cleft region between the head and platform on the 30S subunit (Sharma et al., 2005).

To determine the role of Era in the 30S subunit assembly, we employed quantitative mass spectrometry (qMS), high-resolution cryo-electron microscopy (cryo-EM) and in-cell footprinting to analyze 30S subunit assembly intermediates (30S_Era-depleted_ particles) that accumulated in *Escherichia coli* under Era depletion conditions. In addition, we investigated a potential role of Era in ribosomal quality control and we further explored the functional interplay between Era and another assembly factor, YjeQ (Campbell and Brown, 2008).

Here, we found that in the absence of Era, all of the major 16S rRNA domains fold correctly with the exception of helices 23 and 24 in the platform region, suggesting that maturation of these helices directly or indirectly relies on Era. Notably, our structures indicate that the assembling particles did not stall at the maturation step folding helices 23 and 24. Instead, particles skipped the folding of these two helices and were re-routed in their folding pathway to continue the maturation of other structural motifs. This analysis suggests that assembly of the 30S subunit is not necessarily sequential (from 5’ to 3’) and that at least in the absence of Era, all three major domains of the 16S rRNA can co-mature. Further, we found that treatment of mature 30S subunits with Era destabilizes functionally essential regions of the particle and inhibits binding of YjeQ. Overall, these results suggest that Era binds to the immature 30S subunit independently from YjeQ. The observed ability of Era to induce significant conformational changes in the mature 30S subunit also suggests new roles of this protein in ribosome homeostasis.

## RESULTS

### Era depletion causes accumulation of 30S subunit assembly intermediates

We created a strain that allowed for the controlled expression of Era under an arabinose inducible promoter and we used it to determine how Era depletion impacts the distribution of ribosomal particles and their protein composition in *E. coli* cells. This strain exhibited a growth phenotype consistent with previously characterized Era-depleted strains (Supplemental results and Figure S1). Crude ribosomes were extracted from cells grown under permissive (+arabinose) and restrictive (-arabinose) conditions, layered onto a 10%–30% (w/v) sucrose gradient and subjected to ultracentrifugation. In wild type *E. coli* cells, between 2-5% of the 30S subunits remain typically dissociated instead of forming 70S ribosomes (Leong et al., 2013; Thurlow et al., 2016). Consistent with that data, 3% of the 30S subunits were found dissociated from 50S subunits when the strain was grown under permissive conditions. However, growth in restrictive conditions increased the percentage of free 30S subunits to 72%, significantly decreasing the proportion of 70S ribosomes (Figure 1A; top panel). This percentage was significantly higher than what has been reported in the past for other non-essential assembly factor knock-out strains (*ΔrimM, ΔyjeQ, ΔrbfA*) (Guo et al., 2013; Himeno et al., 2004; Leong et al., 2013; Thurlow et al., 2016) consistent with a more substantial defect in the 30S subunit assembly process.

**Figure 1.**
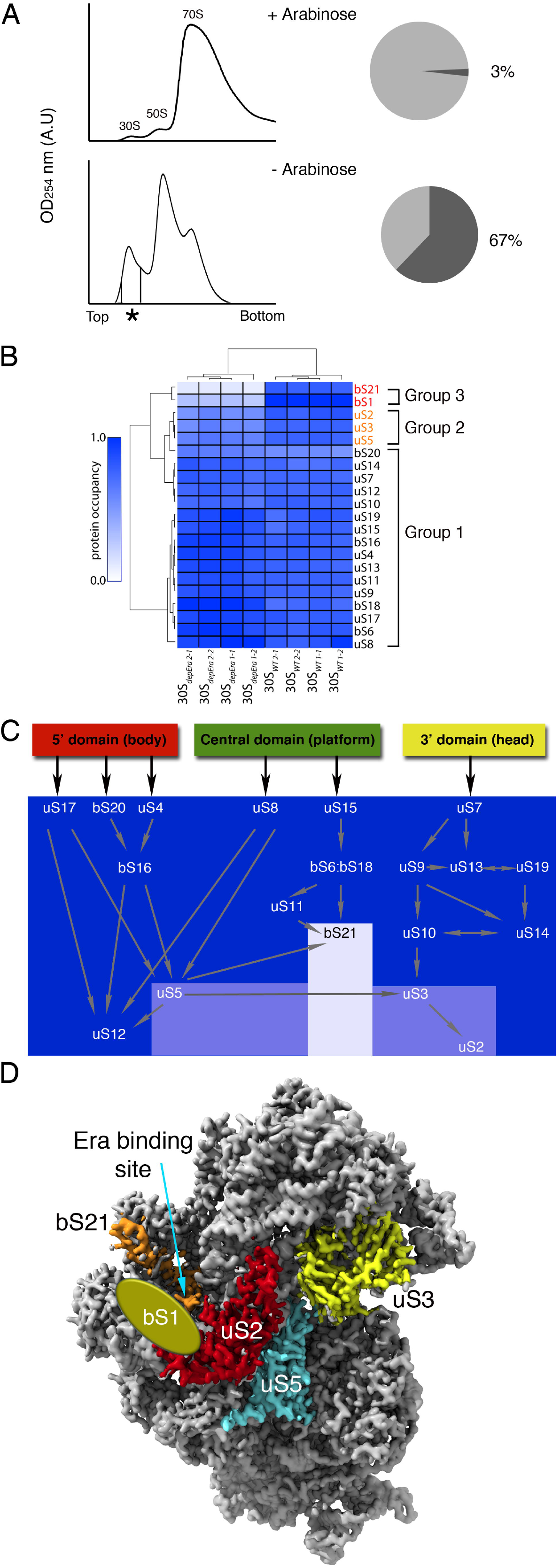
Ribosomal proteins and assembly factors content of the 30S particles in Era depleted cells. (A) Ribosomal profiles of the Era depleted strain under Era expression (+arabinose) and Era depletion (-arabinose) conditions. The asterisk and vertical lines in the bottom gradient indicate the fractions that provided the purified 30S_Era-depleted_ particles for the qMS and cryo-EM experiments. (B) Heatmap showing protein abundance of the 30S particles purified from the parental strain and the Era depleted strain grown under (-arabinose) conditions. Protein abundance is shown relative to a purified 70S particle. Occupancy patterns were hierarchically clustered revealing three groups, which are marked. Proteins in each group are labeled in a different color. Sample replicas were also hierarchically classified and clustered in two groups indicated as 30S_depEra_ (30S_Era-depleted_ particles purified from Era depleted strain grown under (-arabinose) conditions) and 30Swt (30S subunits purified from parental strain). (C) Protein occupancy levels from the qMS analysis of the 30S_Era-depleted_ particles was plotted in the Nomura assembly map of the 30S subunit using the same color coding as in panel A. (D) The r-proteins found to be sub-stoichiometric by qMS are shown in a color different from grey in the structure of the mature 30S subunit. The entire r-protein bS1 is not shown for clarity. The oval shape labeled as bS1 indicates the binding site of the N-terminal region of this protein to the 30S subunit. Ribosomal proteins found on full occupancy in the 30S_Era-depleted_ particles are shown in grey.

Next, we used quantitative mass spectrometry (qMS) to determine the r-protein composition of the 30S particles purified from either parental or Era-depleted strains. Based on their occupancy levels in the 30S_Era-depleted_ particles, r-proteins were divided into 3 groups (Figure 1B and Figure S2). Group 1 included most r-proteins, which were present stoichiometrically in all 30S particles, suggesting that their binding was independent of Era. Conversely, group 3 proteins bS1 and bS21 were completely missing from 30S_Era-depleted_ particles revealing that their binding was severely affected by Era depletion. Finally, group 2 proteins uS2, uS3 and uS5 showed intermediate occupancy levels in 30S_Era-depleted_ particles, suggesting a partial Era-dependence for their binding. We noted that occupancy of bS20 was sub-stoichiometric in small subunit particles derived from either Era-depleted cells or parental cells, consistent with protein dissociation during purification of the subunits.

Painting the Nomura map (Traub and Nomura, 1968, 1969a, b) according to these r-protein groups (Figure 1C) revealed that the protein complement of the 30S_Era-depleted_ particles is consistent with the binding hierarchy defined by the assembly map. Inspection of mature 30S subunit crystal structure (Schuwirth et al., 2005) and cross-linking mass spectrometry data localizing bS1 binding site (Lauber et al., 2012) showed that sub-stoichiometrically bound proteins (group 2 and 3) associate near the Era binding site on the platform region of the particle (Sharma et al., 2005) (Figure 1D). The conspicuous co-localization of these r-proteins suggested that Era may be assisting entry of these proteins.

Taken together this analysis showed that a large proportion of the particles isolated form Era-depleted cells were immature and in the late stages of the assembly process.

### The immature 30S particles accumulating in the Era-depleted strain range from early to late assembly intermediates

To further understand the role of Era in the assembly of the 30S subunit, we determined an ensemble of structures present in the 30S_Era-depleted_ particles using high-throughput, single particle cryo-EM. Our dataset included over 800,000 particle images of purified 30S_Era-depleted_ particles and, after subjecting this dataset to multiple rounds of 3D image classification and initial map determination, we obtained a collection of sixteen cryo-EM maps (Figure 2).

**Figure 2.**
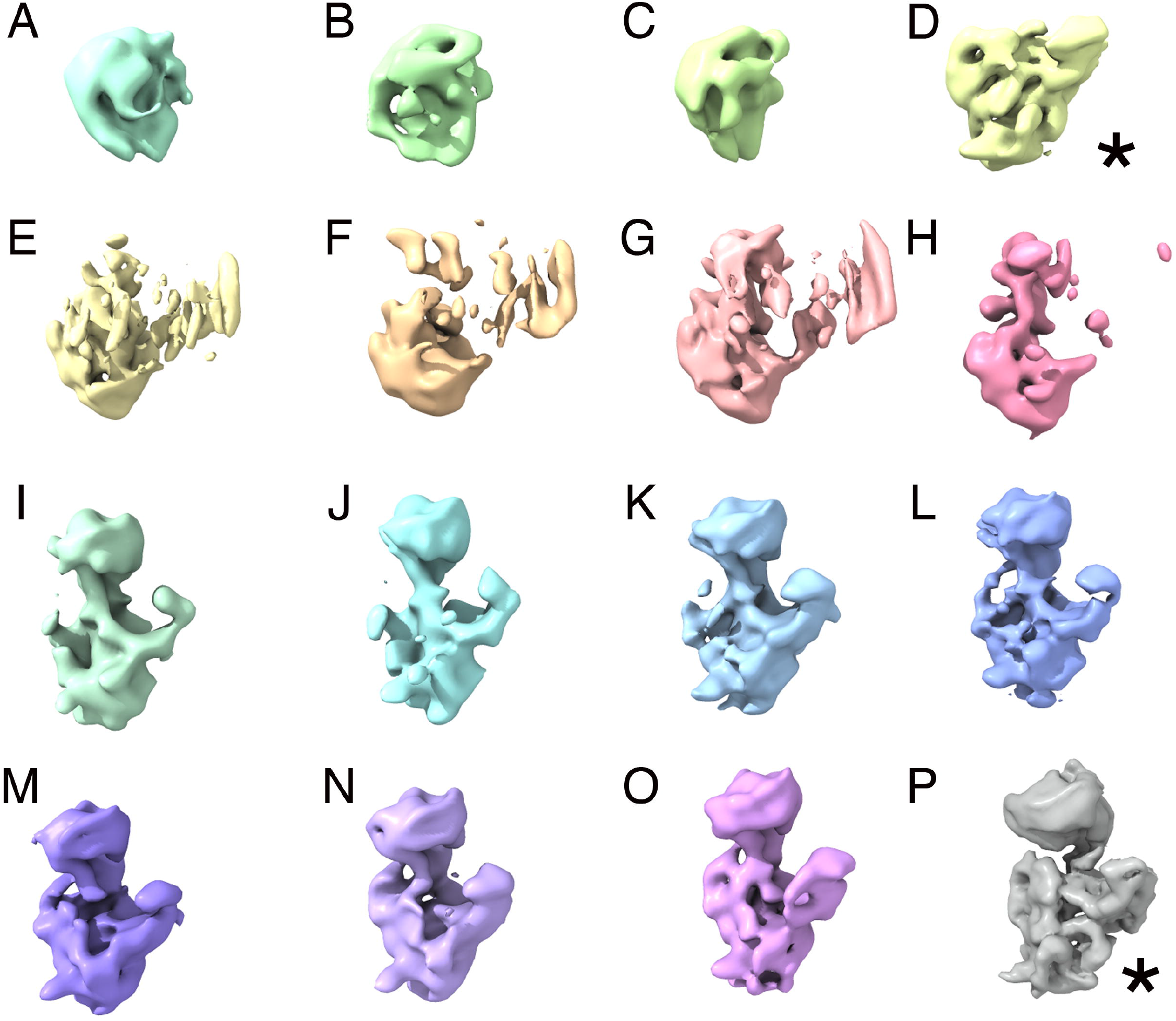
Immature 30S particles accumulating in the Era-depleted strain. (A-P) Cryo-EM maps obtained from a sample containing purified 30S_Era-depleted_ particles using 3D image classification approaches. The most abundant classes are marked by an asterisk. Classes D and P represented the 22% and 50% of the population, respectively. The remaining classes accumulated only a 28% of the particle population.

These sixteen cryo-EM maps provide a direct visualization of the 30S subunit as it assembled (Movie 1). Classes A, B, and C represented particles at very early stages of assembly. These particles exhibited density corresponding to the body of the 30S subunit and lacked platform or head domains. Notably, in these maps the body had not fully matured as rRNA helices could not be readily assigned. Class D contained particles appearing to undergo maturation of the body and platform, but without starting to significantly fold the head. Classes E to H appeared to be folding the three domains simultaneously. The remaining classes (I-P) resembled successively more mature assembly intermediates with body, platform and head domains progressively closer in structure to the mature subunit. No class corresponding to fully mature 30S subunit was observed.

Our structures of isolated body domains (classes A-C), which are composed of the 5’ and central domains are largely in agreement with previous work establishing that rRNA folding occurs 5’ to 3’ (Powers and Noller, 1990; Talkington et al., 2005). Notably however, the observed co-maturation of the body, platform, and head domains in classes E-P indicates that, at least in the absence of Era, rRNA folding is not necessarily sequential and that maturation of all three major domains of the 16S rRNA can occur in concert.

Particles assigned to classes D and P represented the 22% and 50% of the population, respectively and it was possible to obtain high-resolution structures for these assembly intermediates (see section below). All the other remaining classes accumulated only 28% of the particle population and consequently only rendered maps at modest resolution (10-15 Å). However, they still provided sufficient detail to visualize the overall folding of the 16S rRNA domains (Figure 2 and Movie 1).

### Folding of the platform domain of the 30S subunit requires Era

The map for class D refined to 4.8 Å resolution (Figure 3A and Figure S3A). Local resolution analysis showed that the body of the particle is the most defined region, whereas the platform is less well resolved, likely due to intrinsic flexibility caused by the maturation events still ongoing in this domain (Figure S3B). As we were unable to build an atomic model at this resolution, we docked the X-ray structure of a complete 30S subunit (Schuwirth et al., 2005) into this map to visualize how structural elements in class D deviated from the mature structure. This analysis revealed that the density corresponding to the head domain was completely missing in the cryo-EM map of the assembly intermediate whereas density for the rRNA in the body and platform was present. Notably, the body and platform density observed deviated substantially from the tracing of the atomic model of the mature 30S subunit indicating that these regions are still in an immature conformation. The cryo-EM map was also missing density for all the r-proteins in the head, as well as uS2 and uS5 in the body and bS21 in the platform.

**Figure 3.**
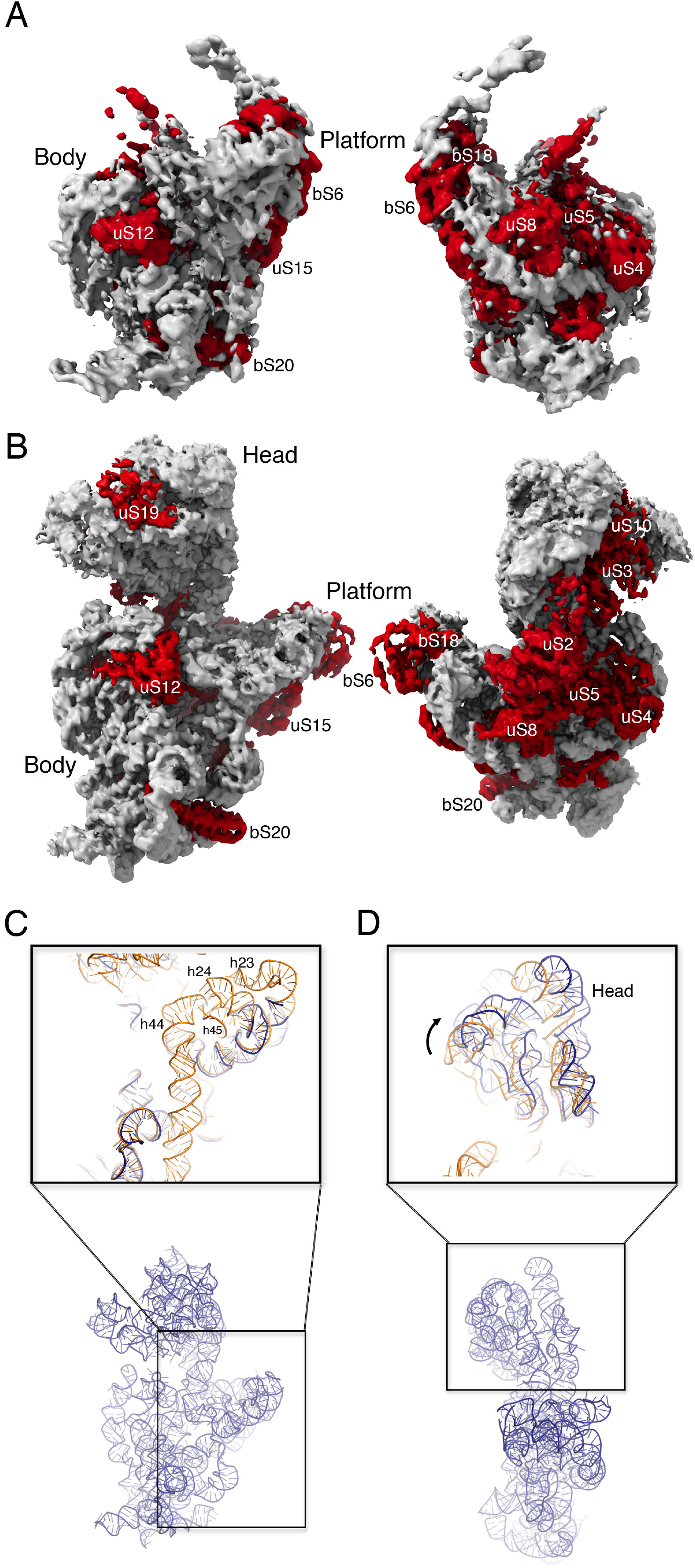
Cryo-EM structures of the most abundant 30S_Era-depleted_ particles. (A) Front and back view of the cryo-EM map of the 30S_Era-depleted_ particle class D. Ribosomal proteins and rRNA in the structure are colored in red and light grey respectively. (B) Similar representation of the cryo-EM map of the 30S_Era-depleted_ particle class P. (C) Zoom-in-view of the 3’ minor and central domains of the atomic model obtained for the 30S_Era-depleted_ particle class P. The area framed with a black square in the overall view of the atomic model (r-proteins removed for clarity) at the bottom is shown enlarged in the top panel. The top panel shows the superimposition of the atomic model for the mature 30S subunit (orange) (PDB ID 4V4Q) and the corresponding region on the model obtained for the 30S_Era-depleted_ particle (dark blue). (D) Zoom-in-view of the 3’ major domain of the atomic model obtained for the 30S_Era-depleted_ particle class P. This region of the model is compared in the top panel with the atomic model of the mature 30S subunit following the same color code as in (C). Ribosomal proteins have been removed for clarity.

The cryo-EM map for class P, the most mature of the assembly intermediates, refined to 3.8 Å resolution (Figure 3B and Figure S3A) with the highest local resolution in the body region of the map. Peripheral domains including the front of the head and platform regions exhibited lower local resolution, again consistent with an intermediate still undergoing maturation. The resolution of this cryo-EM map was sufficient to produce an atomic model from the cryo-EM map (Figure S4), which we used to compare this assembly intermediate to the mature 30S subunit (Figure 3C and Movie 2). The platform and 3’ minor domain exhibited the most striking structural differences. Indeed, in the platform, density for helices 23 and 24 was absent or highly fragmented, and helices 44 and 45 (nucleotides 1397-1534) of the 3’ minor domain were completely invisible (Movie 2). We interpret these fragmented regions of the map as indications that these elements have not fully matured. Finally, all structural elements were present in the head domain, but this entire rigid body was tilted away from the platform (Figure 3D). The remainder of the structure was largely indistinguishable from the mature subunit and exhibited complete density for r-proteins in all four domains except for uS7, uS9, uS13 and uS19 in the head, and uS11 and bS21 in the platform, which each showed a highly fragmented density or lacked density completely (Figure 3B and Movie 2). Combining this structure with our qMS data (Figure 1A), we concluded that proteins bS21 and bS1 are completely absent whereas proteins uS7, uS9, uS11, uS13, and uS19 are likely bound in a flexible manner not easily resolved by cryo-EM. The structural differences of class P with the mature subunit suggests that during the folding of the platform domain, helices 23 and 24 failed to adopt the mature conformation in the absence of Era. However, the assembling particles were able to skip the folding of this motif and re-route their assembly pathway to continue with the maturation of the head domain, which is downstream of helices 23 and 24 (Besancon and Wagner, 1999; Mulder et al., 2010; Sashital et al., 2014; Sykes and Williamson, 2009). Therefore, we concluded that the folding of helices 23 and 24 represents a critical folding barrier during 30S subunit maturation and Era mediates efficient folding of these helices.

### Era expression promotes maturation of helix 23 and 24 in the 30S subunit platform domain

We next used dimethyl sulfate (DMS) footprinting to inspect the structure of the 16S rRNA platform domain with single nucleotide resolution in various Era-limited cells. Specifically, cells were collected before Era depletion (Era pre), after Era depletion (Era depleted) and at various times after Era re-expression (Era+) and were treated with DMS for 0.5 min, 1 min or 2 min (Figure 4A), and the methylation pattern in the 16S central domain was detected by primer extension (Figure 4B and Figure S5). We mapped the 5’ end of the rRNA by primer extension to confirm that the ratio of immature 17S to mature 16S RNA increased upon Era depletion and decreased when Era expression was restored (Figure S6).

**Figure 4.**
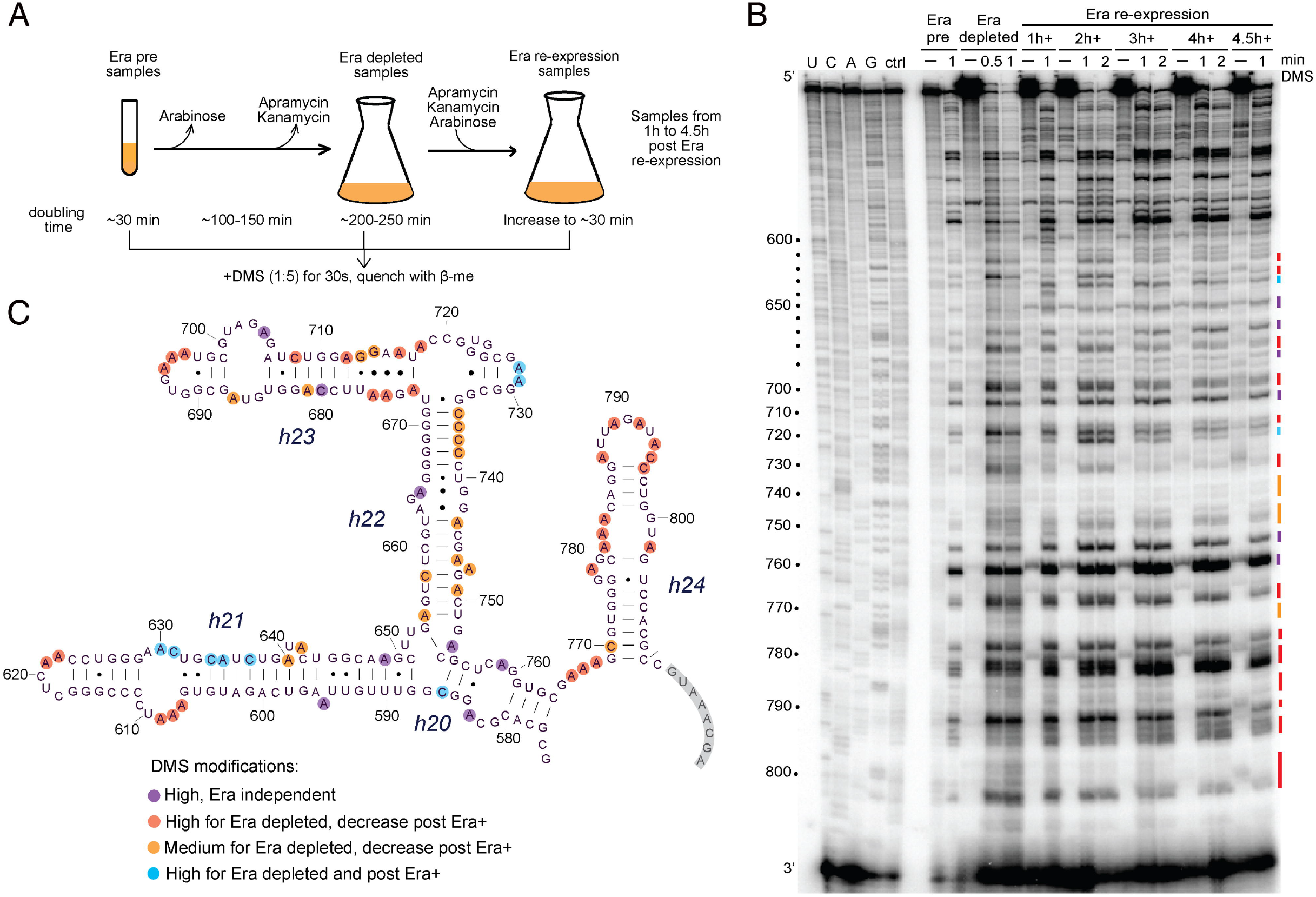
*In vivo* DMS footprinting of Era-dependent changes in the 16S central domain. (A) Scheme of the *in vivo* DMS footprinting experiment after Era depletion and reexpression. (B) DMS modification patterns of different Era samples as detected by extension of primer 812. The Era pre, Era depleted and Era re-expression cells were untreated (–) or modified with DMS for 0.5 min, 1 min or 2 min. DMS modified nucleotides are highlighted with colored bars (right). The nucleotide numbers are indicated to the left of the sequencing ladder, and the unextended primer is marked (3′). (C) DMS-modified nucleotides on the 16S rRNA secondary structure are colored based on the effect of Era as indicated in the key and illustrated in Figure S5. The primer binding location is shaded gray.

We found that helices 23 and 24 in the central domain were highly modified by DMS in Era depleted samples, compared to cells probed before Era depletion (Era pre) or after Era was re-induced (Era+) (red and orange in Figure 4B and 4C). This result is consistent with the cryo-EM maps obtained for the 30S_Era-depleted_ particles purified from cells grown under Era depletion conditions. All of these structures exhibited non-mature conformations of the platform domain and had missing or broken densities for helices 23 and h24. In sum, these results confirm that these two helices were not natively folded in the absence of Era. Likewise, nucleotides in helix 22 also showed increased DMS modification upon Era depletion and enhanced protection when Era was restored, suggesting that Era directly or indirectly stabilizes the structure of helix 22 (orange in Figure 4B and 4C). As density for helix 22 is observed in the class P cryo-EM maps, this chemical probing result implies that helix 22 is likely immature in most of the other classes (classes A to O), which constitute ~50% of all Era-depleted particles (Figure 2). We also observed delayed changes to the DMS modification pattern of helix 21, in which an internal bulge in helix 21 folds only 1 hour after Era re-expression (blue in Figure 4C and Figure S5C). The base of helix 21 is recognized by ribosomal protein uS8, but its tip extends away from the central domain to contact the solvent side of the 30S subunit 5’ domain. This timing suggests that Era enables structural changes at the interface of the 5’ and 3’ domains during 30S assembly, in agreement with previous *in vitro* footprinting results that revealed conformational changes in the region around r-protein uS8 and slow folding of h21 during 30S assembly (Adilakshmi et al., 2008; Calidas and Culver, 2011). Taken together, our in-cell DMS probing results are consistent with the observed cryo-EM structures and highlight Era’s role in assembly of the 30S subunit platform.

### Era binding to the mature 30S subunit destabilizes helix 44 and decoding center

Cells often employ quality control mechanisms to survey nascent ribosomes and ensure their functionality (Karbstein, 2013; Strunk et al., 2011; Strunk et al., 2012). Recently, it was suggested that in addition to its function as an assembly factor, YjeQ may also test the proofreading ability of the 30S subunit (Razi et al., 2017). Thus, to investigate a potential role for Era in ribosome quality control, we assayed Era’s ability to interact with mature ribosomes, and determined the effect of this interaction in the 30S subunit using cryo-EM. We first tested the ability of Era to interact with the mature 30S subunit using a dissociation experiment (Lopez-Alonso et al., 2017). This assay exploits the ability of Era to split 70S ribosomes into 50S and 30S subunits (Sharma et al., 2005). Era was mixed and incubated with purified 70S ribosomes in buffer containing either GDP, GTP, GDPPNP or no nucleotide. Subsequently, ribosomal particles were separated by sucrose density ultracentrifugation (Figure 5A). We found that most of the ribosomes remained as 70S particles after mixing a 5-fold excess of Era in the absence of nucleotide. However, we observed significant 70S dissociation when Era was added in the presence of buffer containing one of the nucleotides. Effective splitting was both nucleotide-and Era-dependent as incubation of 70S particles with GDPPNP alone had no effect.

**Figure 5.**
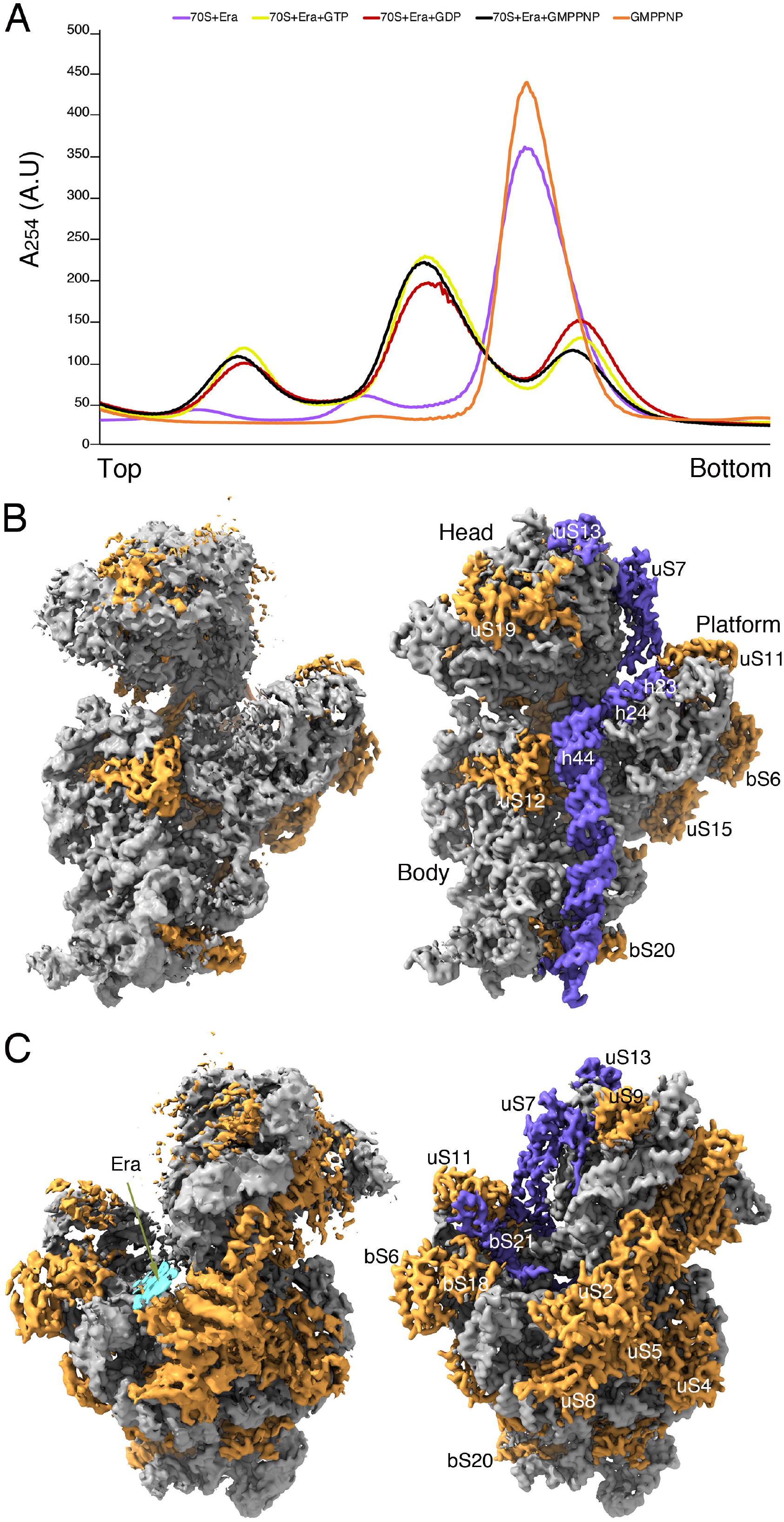
Era dissociates 70S ribosomes and destabilizes helix 44 and decoding center. (A) Ribosome profiles obtained during the 70S dissociation experiments. Plot lines from the reaction containing GDPPNP, GTP, GDP or no nucleotide are shown. All reactions contained a 5-fold excess of Era with respect to the 70S ribosomes. (B) Comparison of the cryo-EM map obtained for the 30S particles treated with Era (left panel) with the structure of the mature 30S subunit (PDB ID 4V4Q) (right panel). The r-RNA and r-proteins are labeled and colored in grey and orange, respectively in both structures. Structural elements absent in the cryo-EM map obtained for the 30S particles treated with Era are colored in blue in the structure of the mature 30S subunit. This panel shows a front view of the structures. (C) Comparison of the back view of the cryo-EM map obtained for the 30S particles treated with Era (left panel) with the structure of the mature 30S subunit (right panel) using the same coloring scheme as in (B). For clarity bS1 is not shown in the mature 30S subunit. The fragmented density potentially representing Era is indicated with an arrow and colored in cyan.

Taking these results into consideration, we mixed mature 30S subunits with Era in a buffer containing GDPPNP and collected images using cryo-EM. Single particle analysis of these images produced a cryo-EM map that refined to 3.9 Å resolution (Figure S7A). Local resolution analysis showed that the highest resolution values were observed in the body domain of the subunit. Peripheral domains including the front of the head and platform regions exhibited lower resolution suggesting higher flexibility in these domains (Figure S7B).

A previous cryo-EM structure showed that in *Thermus thermophilus*, Era bound the mature 30S subunit at the cleft region between platform and head (Sharma et al., 2005). However, the moderate resolution (~13Å) of this structure precluded detailed analysis of any Era-induced structural changes in the 30S subunit. The structure obtained herein (Figure 5B and 5C and Movie 3) revealed significant Era-induced conformational changes in the 3’ minor domain of the mature 30S (Schuwirth et al., 2005). For example, density for helix 44 was completely invisible, and density at the tips of helices 23 and 24 was mildly fragmented (Figure 5B). Density was also lacking for bS1, uS7, bS21, and the N-terminal region of uS13 (Figure 5B and 5C). All other regions of the structure resembled the mature 30S subunit. Surprisingly, we could only observe a small and fragmented density near uS2 in the cleft region between platform and head that could be assigned to Era, suggesting that that unlike *T. thermophilus* Era, *E. coli* Era binds to this region either transiently or in a flexible manner (Figure 5C). In conclusion, this structure suggests that Era has the ability to destabilize functionally essential regions of the mature 30S subunit, including the key elements of the decoding center such as helices 44, 22, and 23.

### Era destabilization of helix 44 blocks binding of YjeQ to the 30S subunit

Previously, Cambell and Brown showed that overexpression of Era suppresses the slow growth phenotype of *ΔyjeQ* cells suggesting a functional interplay between the factors (Campbell and Brown, 2008). Thus, we investigated whether these factors could simultaneously bind the mature 30S, and whether the presence of YjeQ could stabilize Era binding. First, we used microscale thermophoresis (MST) to measure whether pre-binding of Era to the 30S subunit affected the binding of YjeQ to the 30S. Consistent with previous studies (Thurlow et al., 2016), YjeQ exhibited high binding affinity to the 30S subunit with a Kd value of 58.5 ± 47 nM. However, mixing a 10-fold or a 20-fold molar excess of Era with the 30S subunits prior to addition of YjeQ, increased the Kd of the interaction between YjeQ and mature 30S subunits to 1.1 ± 0.6 μM and 2.3 ± 1.2 μM, respectively. These results indicated that binding of Era to the 30S subunit dramatically decreased the affinity of YjeQ for the 30S subunit.

Next, we set up an assembly reaction containing mature 30S subunits plus a molar excess of YjeQ and Era. Single particle analysis produced a cryo-EM map at 3.5 Å resolution (Figure S8A). The highest resolution values were observed in the body domain, whereas the front of the head and platform regions exhibited higher flexibility and produced a map with lower resolution values (Figure S8B).

Previously published structures showed that YjeQ binds to the decoding center and stabilizes helix 44 (Guo et al., 2011; Jomaa et al., 2011b; Lopez-Alonso et al., 2017; Razi et al., 2017). However, we found no additional density in this area of the map that could be assigned to YjeQ and helix 44 was also completely unfolded (Figure 6B). Like the reaction containing mature 30S subunits and Era protein, the cyo-EM map obtained from the reaction containing small subunits and both Era and YjeQ (Movie 4) lacked density for uS7 and the N-terminal region of uS13 (Figure 6B) but showed a defined density for bS21 (Figure 6C). In this case, the map presented a highly fragmented density in the cleft region between the head and platform, which is thought to be the Era binding site to the mature 30S subunit (Sharma et al., 2005). Masked classification on the Era binding region with subtraction of the signal from the rest of the complex did not define this density with sufficient details to unambiguously assign it to Era (Figure 6C). This cryo-EM map allowed us to conclude that the mechanism by which Era decreases the affinity of YjeQ for the mature 30S subunit is by inducing the undocking of helix 44.

**Figure 6.**
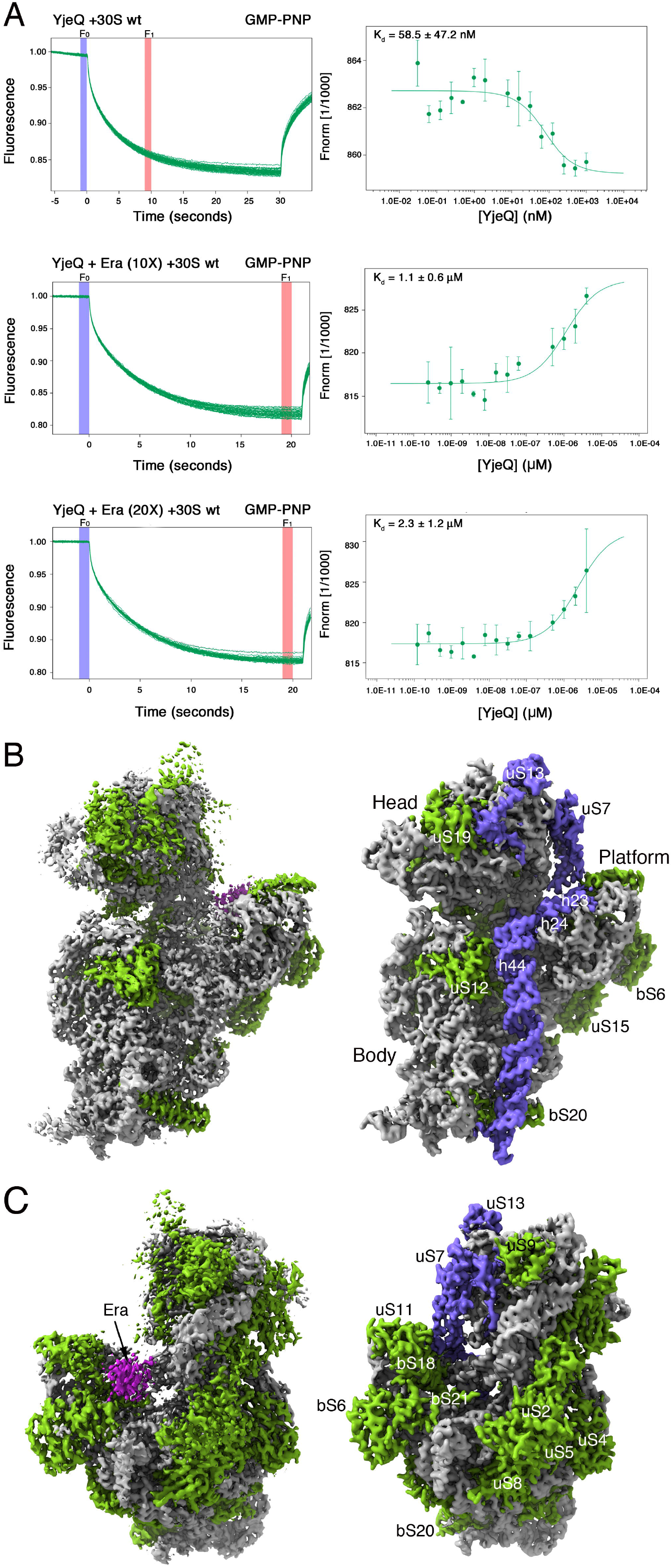
Era blocks binding of YjeQ to the 30S subunit. (A) Analysis of the interaction of YjeQ with the mature 30S subunit by MST. Ribosomal particles were fluorescently labeled and maintained at constant concentration in the three assays. In one assay, the ribosomal subunits were present by themselves and unlabeled YjeQ was titrated into the reaction (top panels). In the other two assays, before YjeQ was added at increasing concentrations, the mature 30S subunits were first mixed with a 10-fold (middle panels) or 20-fold excess of Era (bottom panels). After a short incubation with YjeQ, reactions were loaded into the capillaries and the MST signal of the labeled ribosomal particles (left panels) was measured. Measured changes in the MST response were used to produce curves that plotted the Fnorm (‰) = F1/F0 versus the logarithm of YjeQ concentration. The F1 and F0 regions of the fluorescence time traces used to calculate Fnorm (‰) are indicated in the panel. (B) Front and (C) back views of the cryo-EM map obtained for the 30S particles treated with Era and YjeQ (left panels). Ribosomal proteins and rRNA in the structure are colored in green and light gray respectively. Equivalent views of the mature 30S subunit are shown in the right panels with the structural components displayed using the same color scheme. Structural elements absent in the cryo-EM map obtained for the 30S particles treated with Era and YjeQ are colored in blue in the structure of the mature 30S subunit. For clarity bS1 is not shown in the mature 30S subunit.

## DISCUSSION

### Role of Era in 30S subunit assembly

Prior structural and biochemical analysis of the immature 30S particles that accumulate in *ΔyjeQ* (Jomaa et al., 2011a), *ΔrimM* (Guo et al., 2013; Leong et al., 2013), *ΔrimP* (Sashital et al., 2014) and *ΔyjeQΔrbfA* (Yang et al., 2014) strains have proven powerful in identifying the role of these nonessential assembly factors. Similar approaches have been applied to study the assembly process of the 50S subunit either by genetically removing assembly factors (Ni et al., 2016) or essential r-proteins (Davis et al., 2016). Here, we have extended this methodology to the essential 30S subunit assembly factor Era and employed high-throughput cryo-EM data collection to achieve higher resolution than was possible in prior studies of the 30S.

Previous negative staining EM visualization of 30S particles isolated from wild type *E. coli* and a *ΔrimP* strains (Sashital et al., 2014) and from *in vitro* reconstitution reactions (Mulder et al., 2010) revealed that assembly of the 30S subunit is sequential (from 5’ to 3’) and that folding of the head domain only occurs upon completion of the folding of the 5’ and central domain. In striking difference, our cryo-EM analysis suggests that in the assembly of the 30S subunit, at least in the absence of Era, all three major domains of the 16S rRNA can co-mature. In particular, classes E to H identified within the 30S_Era-depleted_ particle population (Figure 2) appeared to be clearly folding the three domains simultaneously.

Our structures also revealed that most 16S rRNA elements were able to fold even when Era was depleted. Notably, we found that helices 23 and 24 in the platform domain failed to mature upon Era depletion. Given their proximity to the putative Era binding site, we posit that Era is directly involved in the maturation of helices 23 and 24. As described above, helices 23 and 24 are thought to mature early in the assembly process when Era is present (Besancon and Wagner, 1999; Mulder et al., 2010; Sashital et al., 2014; Sykes and Williamson, 2009) and, if ribosome assembly is strictly sequential, one would have expected assembly to stall at this point and the accumulated intermediates to be very immature (Figure 7). Instead, we observe a significant number of late assembly intermediates (classes G to P), implying that particles re-routed their folding pathway to continue the maturation of their other structural motifs (Figure 7) while waiting for maturation of Era-dependent elements.

**Figure 7.**
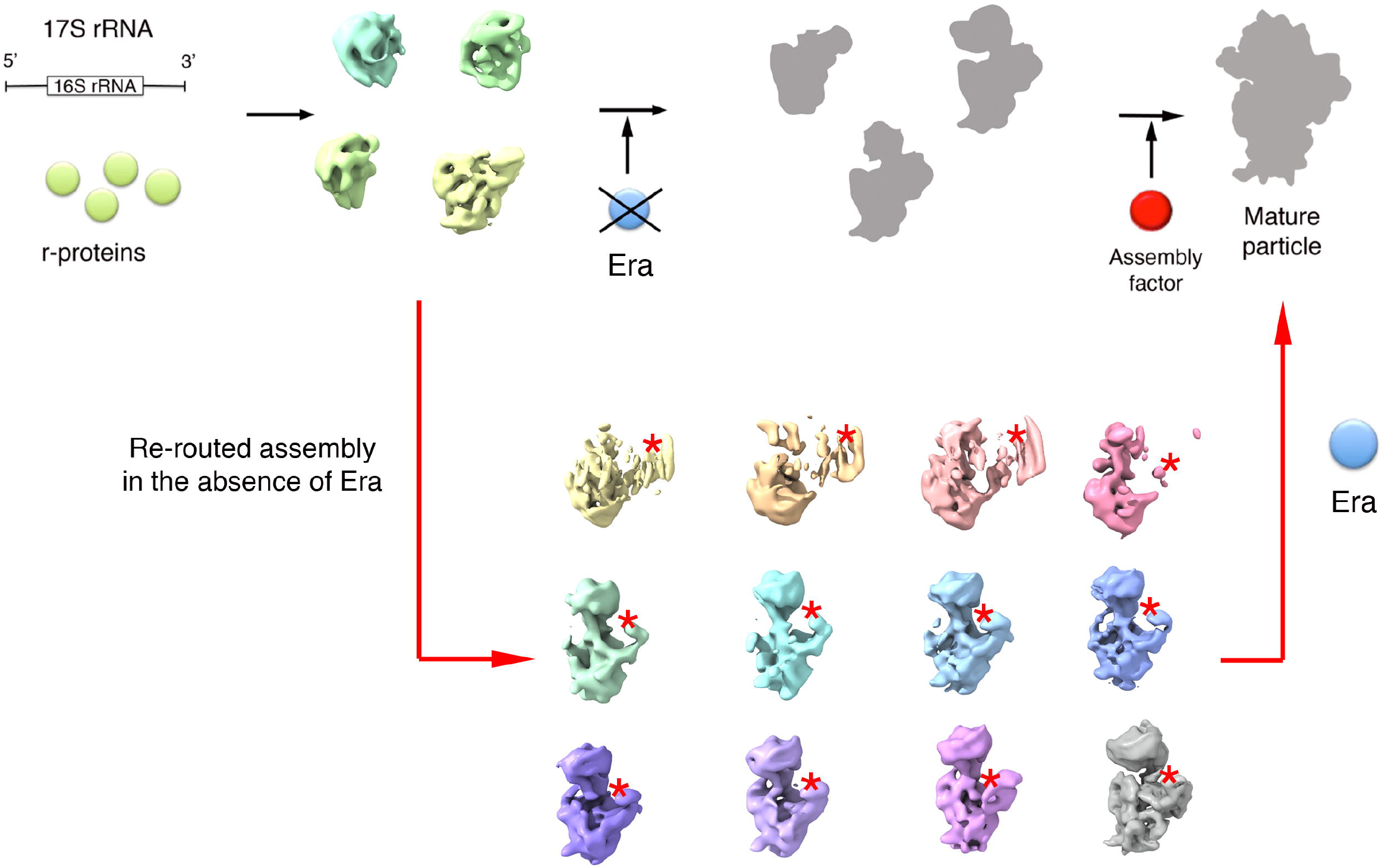
Era depletion causes re-routing of the assembling 30S particles. Diagram summarizing the effect of Era depletion on the assembly of 30S subunits. Cellular depletion of the essential assembly factor Era does not stall the assembly of the 30S subunit, causing accumulation of assembly intermediates upstream of the reaction catalyzed by Era. Rather, in the absence of the Era, the 30S particles are unable to fold helices 23 and 24 in the platform domain, however they re-route their folding pathway and continue the maturation of other structural motifs. During this process, all of the 16S rRNA domains of the 30S subunit fold independently and simultaneously accumulating a variety of assembly intermediates that were characterized in this study using cryo-EM. The red asterisks indicate that helices 23 and 24 still adopt an immature conformation in all of these structures. Only re-introduction of Era in the cells allow these assembly intermediate to reach a mature state. The ribosomal particles shown as grey flat profiles indicate the physiological assembly intermediates that exist in the cell when Era is present at normal concentrations. These particles are capable of folding helices 23 and 24 normally and reach the mature state.

The large percentage (22%) of particles adopting the conformation described by class D probably represent the assembly intermediates unsuccessfully attempting to properly fold these helices, but before finding an alternative assembly pathway to continue their maturation and producing the subsequent classes (classes F to P). This interpretation aligns well with previous studies establishing that assembly of the 30S subunit occurs through multiple parallel assembly pathways (Adilakshmi et al., 2005; Talkington et al., 2005) and recent work illustrating dynamic re-routing of assembly flux through parallel pathways in the large subunit (Davis et al., 2016).

Interestingly, helices 44 and 45 are also immature in each of our structures suggesting that their maturation either directly depends on Era or, instead, requires proper maturation of helices 23 and 24 in the platform domain. We favor the later model as folding of helix 44 and 45 including the decoding center are thought to be final maturation steps before the 30S subunit becomes functionally active and the factors YjeQ and RbfA have previously been implicated in facilitating this step (Clatterbuck Soper et al., 2013; Jomaa et al., 2011a; Lopez-Alonso et al., 2017; Razi et al., 2017).

### Role of Era in ribosome homeostasis

Prior structural work (Sharma et al., 2005; Tu et al., 2009) has also suggested that Era might block premature translation initiation by *immature* small subunit particles. Suggested models include: i) blocking access to the anti-Shine-Dalgarno (SD) sequence near the 3’ end of the 16S rRNA; and ii) preventing the binding of bS1, a r-protein necessary for the SD / anti-SD interaction to the 30S subunit. The 70S dissociation experiments performed herein indicate that Era may also prevent translation initiation by *mature* 30S subunits. Our subsequent structural analysis of the mature 30S particles treated with Era revealed a complete unfolding of helix 44 and reversion of the decoding center to an immature state. As helix 44 contains the b2a inter-subunit bridge that is essential for subunit joining, this Era-treated particle would not be competent for translation.

Reversion of ribosomal structural motifs to an immature state is not unprecedented. Indeed, earlier studies have described this phenomenon upon removal of r-proteins (Jomaa et al., 2014) or binding of the assembly factor YjeQ (Lopez-Alonso et al., 2017). Our analysis demonstrates that Era shares this ability and opens the possibility that cells may utilize these assembly factors to revert mature, active ribosomes into inactive, immature particles. The physiological conditions in which this function of Era or other assembly factors sharing this ability becomes relevant for survival or fitness is still unclear. Possibilities include the participation of Era on active dissociation of hibernating RMF (ribosome modulation factor) HFP (hibernation promoting factor)-100S and RaiA (ribosome-associated inhibitor A)-70S ribosomes that *E. coli* assembles during stationary phase to stop protein synthesis and reduce energy consumption (Prossliner et al., 2018). A recent publication has provided the first insight into GTP-dependent dissociation mechanisms of 100S ribosomes by the widely conserved GTPase HflX (Basu and Yap, 2017). However, deletion of hflX had only a moderate effect on the fraction of dimerized ribosomes, suggesting that HflX is not solely responsible for dissociation and that other protein factors such as Era, could also be involved in the reactivation mechanisms. It is also plausible the cell uses Era and other destabilizing assembly factors to initiate ribosome degradation during starvation. Conditions that increase the formation of ribosomal subunits lead to enhanced degradation, while conditions favoring the presence of intact 70S ribosomes prevent or reduce breakdown (Zundel et al., 2009). Understanding the physiological conditions triggering these potential new functions of Era in ribosome homeostasis remains to be investigated.

### Functional interplay between Era and YjeQ

Prior genetic experiments suggested functional interactions between Era and other assembly factors. For example, overexpression of Era suppressed the ribosome assembly defects in a *ΔrbfA* strain (Inoue et al., 2003), and overexpression of KsgA, which methylates residues in h45, suppressed the cold-sensitive phenotype of Era E200K (Lu and Inouye, 1998). Although these factors are thought to bind at non-overlapping sites on the 30S subunit, (Datta et al., 2007; Sharma et al., 2005), these results suggested that Era could facilitate the function of RbgA or KsgA and potentially substitute for them.

The present study investigates the functional interplay between Era and YjeQ and extends the work of Campbell and Brown (Campbell and Brown, 2008), who reported that Era overexpression increases polysome count and ameliorates the growth defect of a *yjeQ* null strain. YjeQ is thought to act during 30S subunit assembly by aiding in folding of helix 44 and decoding center (Guo et al., 2011; Jomaa et al., 2011a; Jomaa et al., 2011b; Lopez-Alonso et al., 2017; Razi et al., 2017). Our work suggests that Era has the opposite effect on mature ribosomes, as it appears to promote unfolding of helix 44 during the Era-dependent subunit splitting reaction (Figure 6). This result implies that Era binds to the immature 30S subunit independently from YjeQ. However, it remains unclear from this data how Era over-expression compensates for YjeQ function.

### Role of Era in processing and folding of the rRNA

Given that Era recognizes the 1531AUCACCUCC1539 sequence, which is close to the mature 3’ end of 16S rRNA (Ji, 2016; Tu et al., 2009), Era has also been implicated in rRNA processing regulation. In these models, it is suggested that Era binding to this sequence could induce a conformational change in the 17S rRNA that promotes auto-cleavage or exposes the 3’ end of the 17S rRNA to RNases (Inoue et al., 2003). Alternatively, crystallography studies (Ji, 2016; Tu et al., 2009) propose that Era binding actually protects the rRNA from premature cleavage by RNAses during the ribosome assembly process. Our finding that Era depleted cells have higher ratios of 17S rRNA molecules containing the 5’ precursor sequence but not the 3’ sequence (Supplemental results) is consistent with Era playing a protective role by ensuring that the processing of the 3’ end does not occur prematurely. This putative protective role is also consistent with a recent model (Ghosal et al., 2018). proposing that the endoribonuclease YbeY forms a complex with Era to guide accurate RNAse-mediated cleavage of the terminal 33 nucleotides at the 3’ end of the 17S rRNA.

Overall, this study found that folding of helices 23 and 24 in the platform region of the 30S subunit directly or indirectly relies on Era. In the absence of this factor, particles skipped the folding of these two helices and were re-routed in their folding pathway. The assembling particles did not follow a sequential (from 5’ to 3’) folding and all three major domains of the 16S rRNA co-matured. In addition, we found that treatment of mature 30S subunits with Era destabilizes functionally essential regions of the 30S subunit. Further investigations in this newly discovered capacity of Era will unravel the putative role of this essential protein factor in mechanisms of ribosome homeostasis, including the reactivation of hibernating ribosomes or the triggering of ribosome degradation.

## STAR*METHODS

Detailed methods are provided in the online version of this paper.

## SUPPLEMENTAL INFORMATION

Supplemental information includes supplemental results, six figures and one table and can be found with this article online at http:/XXXXXX.

## AUTHOR CONTRIBUTIONS

A.R. designed cryo-EM experiments, performed cryo-EM sample preparation, EM data collection and image analysis. J.D. performed the qMS experiments. Y.H. performed footprinting experiments, D.J. performed MST and 70S dissociation experiments. B.T. obtained the Era depleted strain. K.B. designed and implemented strategies for cryo-EM data collection in the Titan Krios TEM (FEMR-McGill). N.J performed qPCR analysis. J.G and J.V. assisted with designing image processing approaches. A.G. performed model building of the atomic models. R.B, J.W, S.W and J.O. designed and supervised the study. A.R. and J.O wrote the first draft of the manuscript and all authors contributed to editing.

## ACKNOWLEDGEMENTS

We are grateful to Drs. Eric Brown and Jean Philippe Cote for their technical assistance in the production of the Era depleted *E. coli* strain. We thank Drs. Kelly Sears, Mike Strauss and other staff members of the Facility for Electron Microscopy Research (FEMR) at McGill University for help in microscope operation and data collection. We acknowledge Jingyu Sun for providing purified Era for MST experiments and Clara Ortega for assistance with graphic design. This work was supported by grants from the Canadian Institutes of Health Research (PJT-153044) to J.O and NIH grant R01GM110248 from NIGMS to RAB. Titan Krios cryo-EM data were collected at FEMR (McGill). FEMR is supported by the Canadian Foundation for Innovation and Quebec government.

## STAR*METHODS KEY RESOURCES TABLE

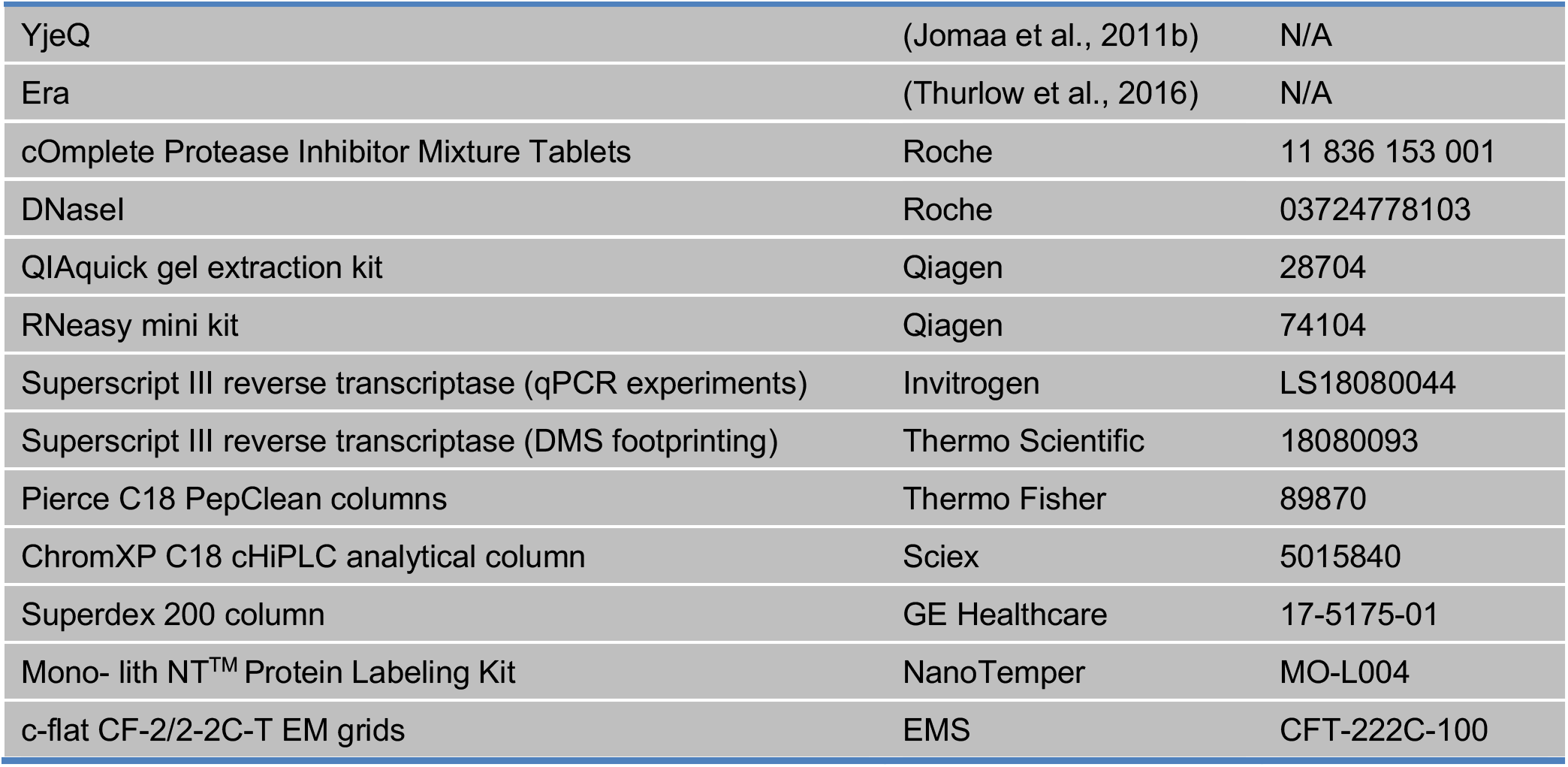
REAGENT or RESOURCE SOURCE IDENTIFIER Chemical, Peptides, and Recombinant Proteins.

Deposited data.

Deposited data
|  |  |  |
| --- | --- | --- |
| Cryo-EM map: 30S<sup>Era-depleted</sup> class D | This paper | EMD: XXXX |
| Cryo-EM map: 30S<sup>Era-depleted</sup> class P | This paper | EMD: XXXX |
| Cryo-EM map: 30S<sup>Era-depleted</sup> class P | This paper | PDB: XXXX |
| Cryo-EM map: Era-treated 30S | This paper | EMD: XXXX |
| Cryo-EM map: Era+YjeQ-treated 30S | This paper | EMD: XXXX |

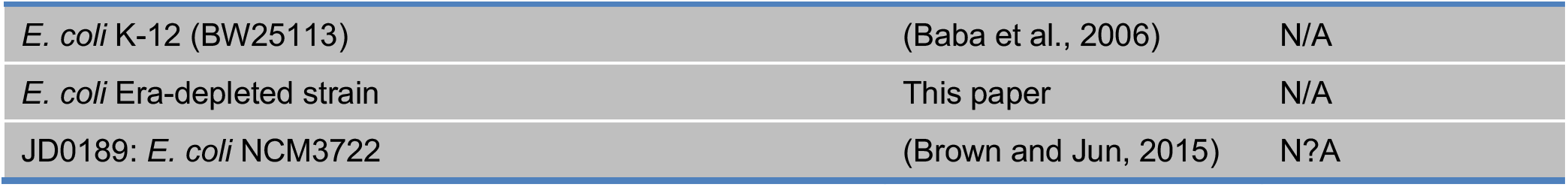
Experimental Models: Organisms/Strains.

Experimental Models: Organisms/Strains
|  |  |  |
| --- | --- | --- |
| *E. coli* K-12 (BW25113) | (Baba et al., 2006) | N/A |
| *E. coli* Era-depleted strain | This paper | N/A |
| JD0189: *E. coli* NCM3722 | (Brown and Jun, 2015) | N/A |

Recombinant DNA.

Recombinant DNA
|  |  |  |
| --- | --- | --- |
| pET15b-*era* | (Thurlow et al., 2016) | N/A |
| pDEST17-*yjeQ* | (Daigle et al., 2002) | N/A |
| pBS-*araBADflankkan* | (Campbell and Brown, 2002) | N/A |
| pSim6 | (Datta et al., 2006) | N/A |
| pBS-*araBADflankerakan* | This paper | N/A |

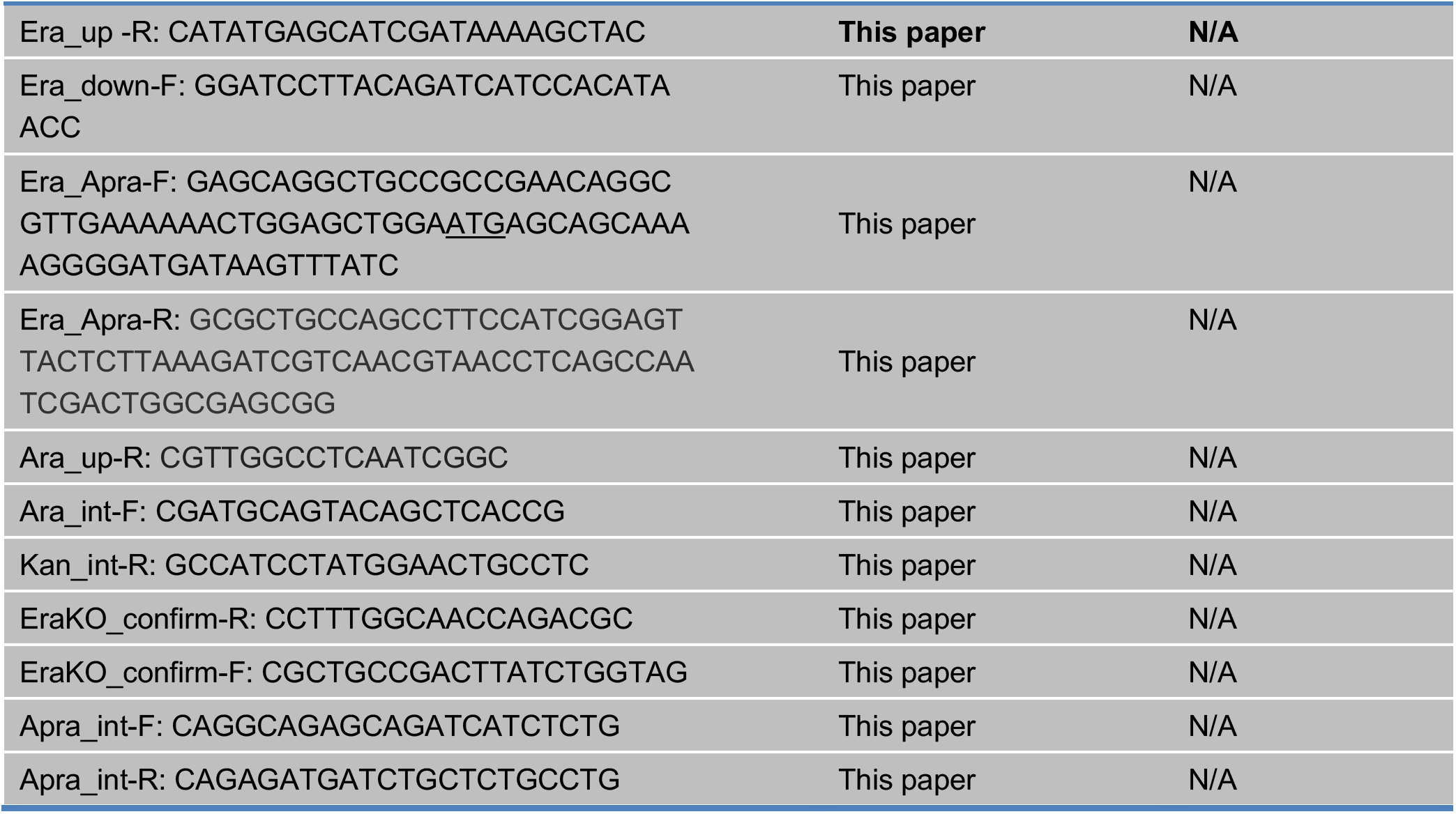
Sequence based reagents. Oligonucleotides used for creation and screening of Era-depleted strain.

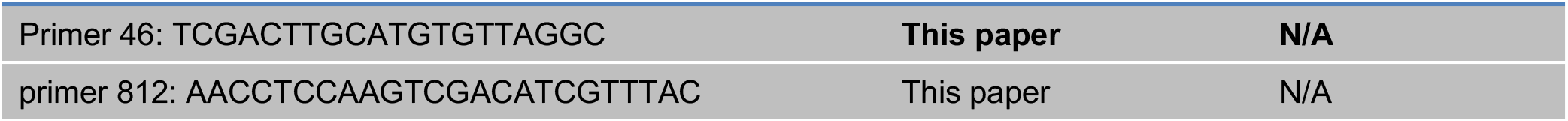
Sequence based reagents. Oligonucleotides used for primer extension.

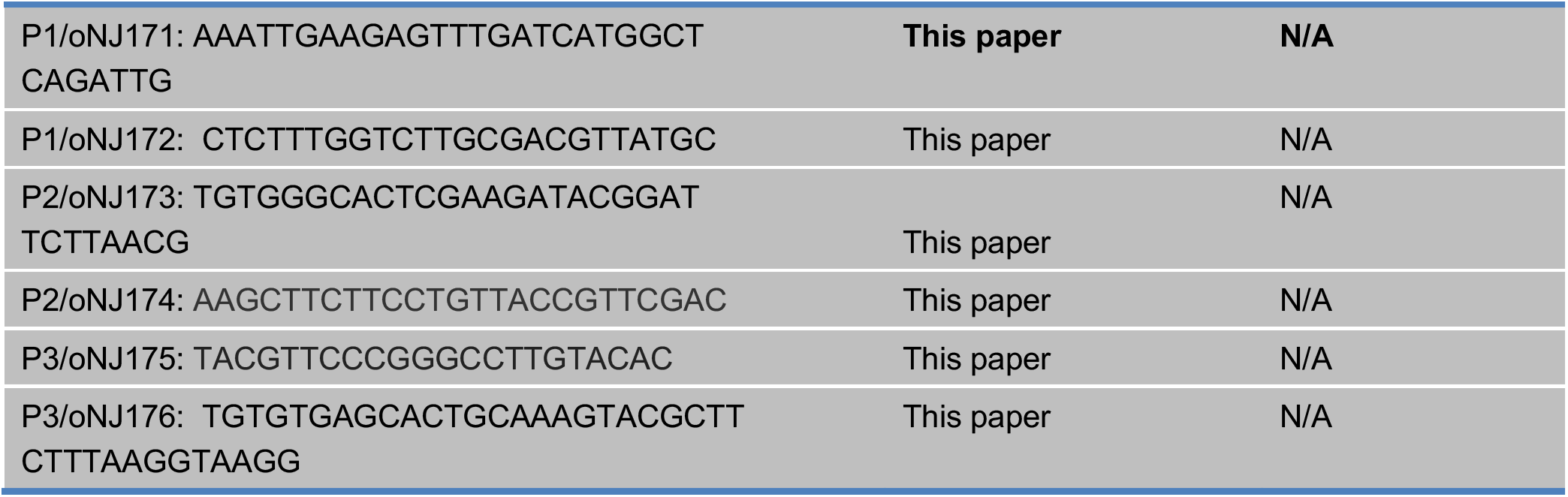
Sequence based reagents. Oligonucleotides used for qPCR.

Software and Algorithms.

Software and Algorithms
|  |  |  |
| --- | --- | --- |
| Comet | (Eng et al., 2013) | N/A |
| PeptideProphet | (Deutsch et al., 2010) | N/A |
| Spectrast | (Lam et al., 2007) | N/A |
| iProphet | (Shteynberg et al., 2011) | N/A |
| Skyline | (MacLean et al., 2010) | N/A |
| NanoTemper MO Affinity Analysis software | NanoTemper | N/A |
| MotionCor2 | (Zheng et al., 2017) | N/A |
| Gctf | (Zhang, 2016) | N/A |
| Relion 2.1 | (Kimanius et al., 2016) | N/A |
| phenix.auto_sharpen | (Terwilliger et al., 2018) | N/A |
| PHENIX software suit | (Afonine et al., 2018) | N/A |
| Coot | (Emsley et al., 2010) | N/A |
| EMRinger | (Barad et al., 2015) | N/A |
| Pymol, Version 1.2r3pre | Schrödinger, LLC | N/A |
| UCSF Chimera and Chimera X | (Pettersen et al., 2004) | N/A |
| ImageQuant | GE Healthcare | N/A |

## CONTACT FOR REAGENT AND RESOURCE SHARING

Further information and requests for resources and reagents should be directed to and will be fulfilled by the Lead Contact, Joaquin Ortega.

## EXPERIMENTAL MODEL AND SUBJECT DETAILS

*E. coli* depletion strain and the strain for overexpression of Era were grown as noted in the STAR Methods section.

## METHOD DETAILS

### Cell strains and protein overexpression clones

The parental *E. coli* K-12 (BW25113) strain was obtained from the Keio collection, a set of *E. coli* K-12 in-frame, single gene knockout mutants (Baba et al., 2006).

The pET15b-era plasmids used for overexpression of Era were produced as described (Thurlow et al., 2016). The pDEST17-yjeQ plasmid used to overexpress YjeQ protein with an amino-terminal His6 tag cleavable by tobacco etch virus (TEV) protease was generated as described previously (Daigle et al., 2002).

### Protein expression and purification

Overexpression and purification of Era (Thurlow et al., 2016) and YjeQ (Razi et al., 2017) proteins was done as previously described.

### Creation of the Era-depleted strain

To generate the essential *era* gene depletion strain in the BW25113 *E. coli* background, we used the previously described pBS-araBADflankkan plasmid (Campbell and Brown, 2002) and the temperature sensitive pSim6 plasmid (Datta et al., 2006) for insertion of *era* at the *araBAD* locus and recombination into the chromosome, respectively. PCR amplification of the *era* gene was performed with the Era_up-R and Era_down-F oligonucleotides using the previously described pET15b-era plasmid (Thurlow et al., 2016) as a template for amplification. The amplified product was cloned into the *PmeI* site of the pBS-araBADflankkan and the resulting plasmid with *era* in the forward orientation with respect to the *araBAD* promoter was sequenced and named “pBS-araBADflankerakan”. The *era* knockout cassette was generated by PCR amplification of pSET152 (Bierman et al., 1992) with the Era_Apra-F and Era_Apra-R oligonucleotides. This generated an approximately 1100 bp product that contained the apramycin^r^ cassette flanked by 50 bp of the upstream and downstream sequences of the native *era* gene.

The precise deletion of *era* was carried out as described by (Datsenko and Wanner, 2000; Link et al., 1997). pBS-araBADflankerakan was double digested with *NotI* and *PsiI* resulting in an approximately 3,200–bp and 2,600-bp product. The digestion reaction was resolved in a 1% agarose gel and the 3.2 kb product containing the *era* gene flanked by homologous regions to the *araBAD* promoter was excised and gel purified using the QIAquick gel extraction kit (Qiagen). Approximately 100 ng of this product was used to transform BW25113-pSim6 competent cells using electroporation and cells were plated on LB-agar containing 50 μg/ml kanamycin at 37 °C overnight to select for integrants at *araBAD* and facilitate curing of the temperature-sensitive pSim6 plasmid. To screen for strains in which the *araBAD* genes had been replaced by *era*, genomic DNA was isolated from the clones and used as a template for PCR amplification with the Era_down-F and Ara_up-R oligonucleotides (Figure S1A; top panel). A second round of PCR screening was performed using the Ara_int-F and Kan_int-R oligonucelotides (Figure S1A; bottom panel). A strain positive for chromosomal integration was selected and named “*araBAD-era*”. The *era* gene deletion protocol was conducted by transforming approximately 100 ng of the *era* knockout cassette using electroporation into *araBAD-era-pSim6* cells followed by plating onto LB-agar containing 50 μg/ml kanamycin and 100 μg/ml apramycin and overnight incubation at 37 °C to select for integrants at the *era* locus and enable curing of the pSim6 plasmid. Genomic DNA was isolated from clones and used a template for PCR amplification using the EraKO_confirm-R and Apra_int-F oligonucleotides (Figure S1B; top panel) and the Apra_int-R and EraKO_confirm-F oligonucleotides (Figure S1B; bottom panel) were used to confirm the deletion of *era*. The resulting strain with genotype *araBAD::era-kan*^r^, *era::apr*^r^ was called “Era-depleted”.

### Strain growth experiments

To obtain the growth curves for the parental and Era-depleted strains, the cultures were grown overnight in LB media at 37 °C with shaking at 225 rpm in an Excella E24 incubator (New Brunswick). In the case of the Era-depleted strain, the LB media was supplemented with 1% arabinose. The overnight cultures were diluted 1/10,000 with LB media to a final volume of 100 μL in a 96-well plate with fresh LB media and the Era-depleted strain was grown in either the presence or absence of 1% arabinose. Cultures were incubated at either 37°C or 15°C with shaking in a Tecan Sunrise^™^ Plate Reader for either 24 or 72 hours, respectively. Culture density was monitored by measuring the optical density at 600 nm (OD_600_) every ten minutes during the time course of the experiment. Optical density was plotted over time (minutes) to show the growth curve for each strain at both 37 °C and 15 °C. The experiment was performed with five replicates.

Dilution plating experiments were performed by inoculating 0.5 mL of a saturated overnight culture in 50 mL of fresh LB media. Cultures were grown at 37°C with shaking until an OD_600_ of 0.2. Then, serial dilutions were made in 10-fold increments, and 5 μL of each dilution was immediately spotted onto LB agar plates with and without 1% arabinose and incubated at 37°C. The plates were incubated until isolated colonies of the parental strain reached ~2 mm in diameter.

### Purification of ribosomal particles from parental and Era-depleted strain

Purification of mature 30S subunits was done from the parental *E. coli* K-12 (BW25113) strain after 1 L of LB media was inoculated with 10 mL of saturated overnight culture and grown to an OD_600_ of 0.2. Cultures were then cooled down to 4 °C and purification of the mature 30S subunits that were used in the cryo-EM experiments was done as previously described (Razi et al., 2017). In the case of the 30S subunits and 70S ribosomes purified under ‘low salt’ conditions for mass spectrometry, we used a previously published protocol (Leong et al., 2013), but all buffers A to F contained only 60 mM NH4Cl. In addition, the salt wash performed with buffer C was omitted.

In the case of the 30S_Era-depleted_ particles and 70S ribosomes from Era-depleted strain, overnight cultures of this strain were grown at 37 °C with shaking at 225 rpm in LB media containing 1% arabinose, 100 μg/ml apramycin and 50 μg/ml kanamycin. To initiate depletion, cells from saturated overnight cultures were pelleted by centrifugation at 12,000g in a microcentrifuge and all arabinose containing media was removed. Cells were then diluted to OD_600_ of 0.02 in LB media containing 100 μg/ml apramycin and 50 μg/ml kanamycin and grown at 37 °C with agitation until they reached a doubling time of ~150 minutes. Cells were not allowed to grow beyond an OD_600_ of 0.2, thus if at this point cells were still exhibiting doubling times <150 min, then the culture was diluted again to OD_600_ = 0.02 into pre-warmed LB media with the same apramycin and kanamycin but without arabinose. Typically, two cycles of growth dilution were required before the cells reached ~150 min doubling time and then cultures were used to inoculate 4 liters of pre-warmed media to an OD_600_ of 0.02 and cells were grown with agitation until they reached an OD_600_ of 0.2. Full depletion of the Era protein was obtained in cells with a doubling time of 225 - 275 min. At that point, cells were harvested for purification of 30S_Era-depleted_ particles and 70S ribosomes. To purify these particles, we used a similar protocol using ultracentrifugation over sucrose gradients as previously described (Leong et al., 2013). However, all buffers A to F contained only 60 mM NH4Cl. In addition, the salt wash performed with buffer C was omitted. This purified 30S_Era-depleted_ particles and 70S ribosomes were used both for cryo-EM and mass spectrometry experiments. The calculation of the proportion of free 30S subunits in the Era-depletion strain from the sucrose gradient was done as previously described (Thurlow et al., 2016).

### Ribosomal RNA Analysis of Era-depleted and parental strains by qPCR

Quantitative real time PCR (qPCR) was carried out using total RNA isolated from the parental and Era-depleted strain (grown under depletion conditions). The rRNA from each sample was purified using the RNeasy mini kit (Qiagen) according to the manufacturer’s protocol. RNA concentration in the sample was then measured by A260, where 1 absorbance unit is equivalent to 40 μg/mL of RNA. cDNA synthesis was carried out using 100ng of random hexamer primers and 10 ng of total RNA with Superscript III reverse transcriptase (Invitrogen) by following manufacturer protocol. cDNA was amplified by primer set P1, P2 and P3. ΔCt values for P2 and P3 oligo sets amplification for both wild type and Era-depleted strains were calculated against total 16S rRNA fragment measured by P1 set of oligos amplification. ΔΔCt was calculated for ΔCt(Era-depleted)-ΔCt(wild type) and fold expression was calculated by 2^(-ΔΔCt)^. Standard deviations were calculated as described in manufacturer protocol of applied biosystem.

### Quantitative mass spectrometry

Performing the qMS analysis of the ribosomal particles purified from the Era depleted and parental strains required to generate ^15^N-labeled isotopic spikes. Whole cell lysates spikes were generated by growing strain JD0189 (*E. coli* NCM3722 (Brown and Jun, 2015)) to OD_600_ 0.4 in ^15^N-labeled minimal media at 37 °C as described previously (Davis et al., 2016). Cells were pelleted and resuspended in 50 mM Tris, 100 mM NH4Cl, 10 mM MgCl2 pH 7.8 to a concentration of 3.0 OD_600_ units/mL, which corresponds to ~0.05 μM ribosomes, assuming 5×10^8^ cells/OD_600_/mL; 20,000 ribosomes/cell. ^15^N-labeled 70S particles spikes were generated by growing strain JD0189 to OD_600_ 0.7 in ^15^N-labeled media as described above. Cells were lysed, and 70S particles were purified on a 10%-40% sucrose gradient as described (Davis et al., 2016). Fractions bearing monosomes or polysomes were pooled and saved at −80C at 0.055 μM.

To generate the tryptic peptides particles were isolated as described above and concentrated to 0.5 μM. To each sample (10 pmols), an equimolar ^15^N-labeled spike sample (either purified ribosomal particles or lysate bearing ~10 pmols of ribosomes) was added, mixed, and trichloroacetic acid (TCA) was added to 13% final concentration. After overnight precipitation at 4 °C, pellets were sequentially washed with 10% TCA, 100% Acetone and were dried before they were resuspended in 40 μL buffer B (100 mM NH4CO3, 5% acetonitrile, 5 mM dithiothreitol) as described (Jomaa et al., 2014). After reduction for 10 minutes at 65 °C, cysteines were alkylated by addition of iodoacetamide to 10 mM and incubation at 30 °C for 30 mins. Then, 0.2 μg trypsin was added, and protein digestion proceeded at 37 °C overnight. Tryptic peptides were purified on C18 PepClean columns, and ~1 μg of peptides was mixed with 500 fmol iRT retention time standards (Pierce) and injected onto the LC/MS.

To perform the qMS peptides were trapped on a ChomXP C18 cHiPLC column (Sciex) and resolved on a 15 cm ChromXP C18 cHiPLC analytical column using a 120 minute 5-35% linear acetonitrile gradient with a flowrate of 300 nL/min. Data was acquired in replicate with either data-dependent acquisitions or in SWATH data-independent acquisition (DIA) mode. Briefly, data-dependent acquisitions included one MS1 scan (200 msec accumulation time) followed by 30 MS2 scans (100 msec accumulation time). The SWATH DIA method consisted of one MS1 scan (200 msec accumulation time) followed by 32 MS2 scans (25 m/z wide, 100 msec accumulation time) covering the range 400-1200.

To analyze the abundance of the r-proteins in the ribosomal particles, DDA datasets were searched with comet (Eng et al., 2013), peptide-spectra matches (PSM) were scored with PeptideProphet (Deutsch et al., 2010) and scored PSMs were combined with iProphet (Shteynberg et al., 2011). A consensus spectral library was generated using Spectrast (Lam et al., 2007). Using this spectral library, a target list bearing ^14^N- and ^15^N-labeled precursor and product ions corresponding to ribosomal proteins of interest was generated using Skyline (MacLean et al., 2010), and ion chromatograms were extracted in regions of the chromatogram near the predicted elution time as determined using the iRT standards (Escher et al., 2012). Extracted ion chromatograms were filtered for interfering ions at both the MS1 and MS2 level, and relative peptide abundance was calculated as ^14^N_total_area/^15^N_total_area. For each sample and for each protein, the median peptide abundance ratio was then calculated, and these values were clustered across both samples and proteins using the ward linkage method and the Euclidean distance metric (Sturn et al., 2002). Non-ribosomal proteins were analyzed as described using a spectral library focused on assembly co-factors (Davis, Williamson Manuscript in preparation) (Davis et al., 2016).

### *In vivo* DMS footprinting experiments

To prepare the cells for in vivo DMS footprinting, Era-depleted cells *were* grown overnight in LB supplemented with apramycin (100 μg/mL), kanamycin (50 μg /mL) and 1% arabinose at 37 °C with shaking at 200 rpm. A 10-mL aliquot of this overnight culture was collected as the ‘Era pre’ sample to be used for *in vivo* DMS footprinting (see below). Fresh LB medium (150 mL) with apramycin (100 μg/mL) and kanamycin (50 μg/mL) was inoculated with 150 μL overnight culture to achieve an OD_600_ = 0.02. The culture was grown at 37 °C until the doubling time reached 100-150 minutes. This culture was then used to inoculate fresh LB media without antibiotic (initial OD_600_ was 0.02). The culture was grown until the doubling time reached 200-250 minutes, when two 50 mL aliquots were removed as the Era depleted (Era-) samples. A portion of the Era depleted culture was sub-cultured to and OD_600_ of 0.02 into LB media with apramycin (100 μg/mL), kanamycin (50 μg/mL) and 1% arabinose to induce Era expression, and this time was recorded as the start of Era re-expression. 50 mL of samples were collected for analysis and DMS footprinting at 1, 2, 3, 4 and 4.5 h after Era re-expression. As a control for second site suppressors, a portion of the Era-culture was rediluted to an OD_600_ of 0.02 in LB without antibiotics or arabinose. The cultures containing arabinose had a shorter doubling time than the control sample lacking arabinose. During the entire procedure, the growth of all cultures was measured every hour to avoid exceeding an OD_600_ = 0.2, 0.2, at which point cultures were diluted to OD_600_ = 0.02 with fresh medium.

The *in vivo* DMS footprinting was carried out on 50-mL cultures, OD_600_ = 0.2, by adding 5 mL 20% (v/v) dimethyl sulfate in ethanol at room temperature for 30 s. The methylation reaction was quenched with 25 mL of ice-cold 1.4 M β-mercaptoethanol and 25 mL of water-saturated isoamyl alcohol for 30 s on ice. The cells were collected at 4,000 rpm (3,315 x *g)* for 10 min using a JS-5.3 rotor (Beckman Coulter). The cell pellets were resuspended twice in the same volume of 0.7 M β-mercaptoethanol and harvested by centrifugation after each step. The washed pellets were stored at −80 °C prior to RNA extraction.

Total RNA was extracted from each cell pellet using the RNeasy mini-prep (Qiagen), and the final RNA concentration was determined from the UV absorption. The DMS modification pattern was analyzed by reverse transcription as previously described (Hao et al., 2018; Hao and Kieft, 2014). For each reaction, 2 μg total RNA was annealed with 2 pmol (30,000 cpm) ^32^P-labeled primer (primer 46 or primer 812) at 65 °C for 3 min before extension with Superscript III (Thermo Scientific) for 60 min at 55 °C. Sequencing lanes were normalized to the total intensity in each lane, excluding the unextended primer and the full-length cDNA, using ImageQuant software. This was followed by background subtraction of the control lanes (– DMS) in Microsoft Excel.

### 70S dissociation experiments

The parental *E. coli* K-12 strain (BW25113) was used for purification of the 70S ribosomes used in these experiments. Typically, 3 L of LB media were inoculated with 10 mL each of saturated overnight culture and grown to an OD_600_ of 0.6. Cells were harvested by centrifugation at 3,700 *g* for 15 min. Cell pellets were resuspended in 7 mL of buffer A (20 mM TrisHCl at pH 7.5, 10 mM magnesium acetate, 100 mM NH4Cl, 0.5 mM EDTA, 3 mM 2-mercaptoethanol, and a protease inhibitor mixture (cOmplete Protease Inhibitor Mixture Tablets; Roche) and DNaseI (Roche). Each of the subsequent steps was performed at 4 °C. The cell suspension was passed through a French pressure cell at 1,400 kg/cm^2^ three consecutive times to lyse the cells. The lysate was spun at 59,000*g* for 30 min to clear cell debris. Resulting supernatant was layered over a sucrose cushion of equal volume composed of 30% sucrose in buffer B (20 mM TrisHCl pH 7.5, 10 mM magnesium acetate, 500 mM NH4Cl, 0.5 mM EDTA, and 3 mM 2-mecaptoethanol), and then spun down for 4.5 h at 321,000g. The pellet was resuspended in buffer C containing 10 mM TrisHCl pH 7.5, 10 mM magnesium acetate, 500 mM NH4Cl, 0.5 mM EDTA and 3 mM 2-mecaptoethanol and then spun for 16 h at 100,000g. The washed ribosome pellet was resuspended in buffer E containing 10 mM Tris·HCl pH 7.5, 10 mM magnesium acetate, 60 mM NH4Cl, and 3 mM 2-mecaptoethanol, which caused subunits to be associated. Approximately 120 A260 units of resuspended crude ribosomes were then applied to 34 mL of 10–30% (wt/vol) sucrose gradients prepared with buffer E. The gradients were centrifuged for 16 h at 40,000g on a Beckman Coulter SW32 Ti rotor. Gradients were fractionated using a Brandel fractionator apparatus and an AKTA Prime FPLC system (GE Healthcare). The elution profile was monitored by UV absorbance at A254, and fractions corresponding to the 70S subunit peak were pooled and spin down for another 4.5 h at 321,000g on a Beckman SW32 Ti rotor. The pellet was resuspended in Buffer E.

To perform the 70S dissociation experiment purified Era was first passed through a Superdex 200 size-exclusion column (GE Healthcare) equilibrated with buffer containing 50mM TrisHCl pH 8.0. To assemble the dissociation reaction 1uM 70S ribosomes were incubated with either 2- or 5-fold molar excess of Era. Reactions had a total volume of in a 120 μL and were assembled in buffer containing 20mM TrisHCl pH 7.5, 20mM magnesium acetate, 30mM potassium chloride, 4mM 2-mecaptoethanol. Where indicated we added up to 1 mM of nucleotide (GTP/GDP/GDPPNP). After 15-minute incubation at 37°C each reaction was loaded onto 34 mL 10-30% (wt/vol) sucrose gradients prepared with buffer E and centrifuged for 16 h at 40,000g on a Beckman Coulter SW32 Ti rotor. Gradients were fractionated using a Brandel fractionator apparatus and an AKTA Prime FPLC system (GE Healthcare). The elution profile was monitored by UV absorbance at A254.

### Microscale thermophoresis

Mature 30S subunits were fluorescently labeled as described (Thurlow et al., 2016). Prior to each MST experiment, all samples were centrifuged at 14000g for 10 min to remove any aggregates. All reaction and titrations series were prepared in MST buffer containing 20 mM Tris-HCl pH 7.5, 150 mM NaCl, 10 mM MgCl2, 1 mM DTT, 0.05% Tween-20, 0.4 mg/ml BSA and were incubated in in 0.5 ml Protein LoBind eppendorf tubes. Prior to performing the MST experiment, 0.5 μM fluorescently labeled mature 30S subunits were incubated with either 5 μM or 10 μM Era. GDPPNP was added to a final concentration of 1mM and the assembly reaction was incubated for 15 minutes at room temperature. A 1:1 serial dilution of non-labelled YjeQ was then prepared in MST buffer starting at 4 μM. The 30S+Era reaction mixtures were then diluted with MST buffer containing 1mM GDPPNP until the 30S subunits reached a concentration of 40nM. For the titration assay, ten microliters of the corresponding serial dilution of the non-labeled YjeQ was mixed with 10 μl of the fluorescently labeled 30S subunit pre-bound with Era (or without Era for the negative control). Titration reactions were then incubated for 10 min at room temperature loaded into premium capillaries (NanoTemper Technologies) and MST analysis was performed using the Monolith NT.115 (NanoTemper Technologies) at ambient temperature. An LED power of 100% and an MST power of 20–60% were used during MST readings. The resulting binding curves were obtained by plotting the normalized fluorescence (Fnorm (‰) = F1 /F0) versus the logarithm of YjeQ concentration. Kds were calculated using the NanoTemper MO Affinity Analysis software. Experiments were performed in triplicates.

### Cryo-electron microscopy

The 30S_Era-depleted_ particles were prepared for imaging with cryo-EM by preparing a dilution of the particles in buffer E (10mM Tris–HCl at pH 7.5, 10mM magnesium acetate, 60mM NH4Cl and 3mM 2-mercaptoethanol) to a concentration of 30nM that was applied directly to the grid and immediately right after the dilution step. The reactions containing 30S and Era for cryo-EM experiments were prepared as followed. A 20 μL reaction was first prepared in buffer E containing mature 30S subunits (0.5 μM) and Era protein (20 μM). GDPPNP was added to this reaction to a concentration of 2 mM. After the reaction was incubated at 37 °C for 15 min, the reaction was diluted 5-fold in buffer E containing GDPPNP at 2 mM and free Era protein at 4 μM and immediately applied to the EM grid. Finally, in the reaction containing mature 30S, Era and YjeQ for cryo-EM experiments, these components were added to final concentrations of 2 μM, 20 μM and 3.5 μM, respectively. GDPPNP was added to this reaction to a concentration of 2 mM. After the reaction was incubated at 37 °C for 15 min, the assembly reaction was diluted 10-fold in buffer E containing GDPPNP at 2 mM and immediately applied to the EM grid.

In all samples a volume of 3.6 μl of the diluted sample was applied to holey carbon grids (c-flat CF-2/2-2C-T) with an additional layer of continuous thin carbon (5-10nm). Before the sample was applied, grids were glow discharged in air at 5 mA for 15 seconds. Vitrification of samples was performed in a Vitrobot (Thermo Fisher Scientific Inc.) by blotting the grids once for 3 seconds and with a blot force +1 before they were plunged into liquid ethane. The Vitrobot was set at 25 °C and 100% relative humidity.

Automated data acquisition was performed using EPU software at FEMR-McGill using a Titan Krios microscope at 300 kV equipped with a Falcon II direct electron detector (Thermo Fisher Scientific Inc.). Movies for the 30S_Era-depleted_ particles, Era-treated 30S particles and Era+YjeQ-treated 30S particles were collected with a total dose of 35, 46 and 28 e^-^/Å^2^, respectively. All datasets were collected as movies with seven frames acquired in 1 second exposure at a magnification of 75,000x, producing images with a calibrated pixel size of 1.073 Å. The nominal defocus range used during data collection was between −1.25 to −2.75 μm.

### Image processing

Motion correction and contrast transfer function (CTF) estimation for each collected movie was done with MotionCor2 (Zheng et al., 2017) and Gctf (Zhang, 2016) programs, respectively. From here all processing was done with Relion 2.1 program (Kimanius et al., 2016). Particle images after the autopicking process were subjected to reference-free 2D classification to produce a ‘clean’ dataset. The selected particles were subsequently used for hierarchical 3D classification approaches. The initial 3D reference used for these classifications was either a 60 Å low pass filtered map of the mature 30S subunit, the intermediate cryo-EM maps obtained during classification or maps obtained by *ab initio* approaches. The *ab initio* methodologies we used to generate these initial maps were random sample consensus (RANSAC) as implemented in Scipion (Vargas et al., 2014) and common lines as implemented in EMAN2 (Tang et al., 2007). In the case of the 3D classifications of the datasets for the 30S_Era-depleted_ particles and the Era-treated 30S particles no mask was used. However, masks created from the mature structure or intermediate maps obtained during the classification were used in the case of the Era+YjeQ-treated 30S particles. All masks were obtained with ‘relion_mask_create’ command using an initial threshold for binarization of 0.01, extension of the binary mask by two pixels and creating a soft-edges with a width of two pixels. Classes obtained at the end of each classification approach were inspected visually and those deemed to represent the same structures were grouped for refinement. A soft-mask was applied in all refinements.

To improve the details on the density potentially representing Era in the Era-treated 30S particles and Era+YjeQ-treated 30S particles, the corresponding datasets were subjected to focus classification with subtraction of the residual signal using Relion (Scheres, 2012) following an approach previously described (Bai et al., 2015).

Sharpening of the cryo-EM maps was done by applying a negative B-factor estimated using automated procedures (Rosenthal and Henderson, 2003). Local resolution analysis of the structures was also done using Relion procedures (Kimanius et al., 2016).

### Map analysis and Atomic model building

Visualization of the cryo-EM maps was done using Chimera (Pettersen et al., 2004). Before initiating high-resolution model building of the cryo-EM map obtained for the 30S_Era-depleted_ particle (class P), we maximize detail and connectivity of this map using the automatic sharpening tool phenix.auto_sharpen (Terwilliger et al., 2018), which is available as part of the PHENIX software suite (Adams et al., 2010). The starting point for the modeling of this map was the atomic model of the mature 30S subunit (PDB ID 2AVY). This initial model was first fit into the density as a rigid body using Chimera (Pettersen et al., 2004) and the final model was built with successive rounds of real space refinement in Phenix (Afonine et al., 2018) and manual model building in Coot (Emsley and Cowtan, 2004; Emsley et al., 2010). Areas of the map not exhibiting corresponding density were deleted and not incorporated into the models.

Figures for this manuscript were prepared using Pymol program (The PyMOL Molecular Graphics System, Version 1.2r3pre, Schrödinger, LLC), UCSF Chimera and Chimera X (Pettersen et al., 2004).

## QUANTIFICATION AND STATISTICAL ANALYSIS

Quantitation of mass spectrometry data detailed above. Statistical values, where calculated, can be found in the Figure legends.

## DATA AND SOFTWARE AVAILABIILITY

All maps are deposited at EMDB as noted in the Key Resources Table.

## REFERENCES

Adams, P.D., Afonine, P.V., Bunkoczi, G., Chen, V.B., Davis, I.W., Echols, N., Headd, J.J., Hung, L.W., Kapral, G.J., Grosse-Kunstleve, R.W., et al. (2010). PHENIX: a comprehensive Python-based system for macromolecular structure solution. Acta Crystallogr D Biol Crystallogr 66, 213–221.

Adilakshmi, T., Bellur, D.L., and Woodson, S.A. (2008). Concurrent nucleation of 16S folding and induced fit in 30S ribosome assembly. Nature 455, 1268–1272.

Adilakshmi, T., Ramaswamy, P., and Woodson, S.A. (2005). Protein-independent folding pathway of the 16S rRNA 5’ domain. J Mol Biol 351, 508–519.

Afonine, P.V., Poon, B.K., Read, R.J., Sobolev, O.V., Terwilliger, T.C., Urzhumtsev, A., and Adams, P.D. (2018). Real-space refinement in PHENIX for cryo-EM and crystallography. Acta Crystallogr D Struct Biol 74, 531–544.

Anderson, P.E., Matsunaga, J., Simons, E.L., and Simons, R.W. (1996). Structure and regulation of the Salmonella typhimurium rnc-era-recO operon. Biochimie 78, 1025–1034.

Baba, T., Ara, T., Hasegawa, M., Takai, Y., Okumura, Y., Baba, M., Datsenko, K.A., Tomita, M., Wanner, B.L., and Mori, H. (2006). Construction of Escherichia coli K-12 in-frame, single-gene knockout mutants: the Keio collection. Mol Syst Biol 2, 2006 0008.

Bai, X.C., Rajendra, E., Yang, G., Shi, Y., and Scheres, S.H. (2015). Sampling the conformational space of the catalytic subunit of human gamma-secretase. eLife 4, e11182.

Barad, B.A., Echols, N., Wang, R.Y., Cheng, Y., DiMaio, F., Adams, P.D., and Fraser, J.S. (2015). EMRinger: side chain-directed model and map validation for 3D cryo-electron microscopy. Nat Methods 12, 943–946.

Basu, A., and Yap, M.N. (2017). Disassembly of the Staphylococcus aureus hibernating 100S ribosome by an evolutionarily conserved GTPase. Proc Natl Acad Sci U S A 114, E8165–E8173.

Besancon, W., and Wagner, R. (1999). Characterization of transient RNA-RNA interactions important for the facilitated structure formation of bacterial ribosomal 16S RNA. Nucleic Acids Res 27, 4353–4362.

Bierman, M., Logan, R., O’Brien, K., Seno, E.T., Rao, R.N., and Schoner, B.E. (1992). Plasmid cloning vectors for the conjugal transfer of DNA from Escherichia coli to Streptomyces spp. Gene 116, 43–49.

Brown, S.D., and Jun, S. (2015). Complete Genome Sequence of Escherichia coli NCM3722. Genome Announc 3.

Calidas, D., and Culver, G.M. (2011). Interdependencies govern multidomain architecture in ribosomal small subunit assembly. RNA 17, 263–277.

Campbell, T.L., and Brown, E.D. (2002). Characterization of the depletion of 2-C-methyl-D-erythritol-2,4-cyclodiphosphate synthase in Escherichia coli and Bacillus subtilis. J Bacteriol 184, 5609–5618.

Campbell, T.L., and Brown, E.D. (2008). Genetic interaction screens with ordered overexpression and deletion clone sets implicate the Escherichia coli GTPase YjeQ in late ribosome biogenesis. J Bacteriol 190, 2537–2545.

Chen, X., Court, D.L., and Ji, X. (1999). Crystal structure of ERA: a GTPase-dependent cell cycle regulator containing an RNA binding motif. Proc Natl Acad Sci U S A 96, 8396–8401.

Clatterbuck Soper, S.F., Dator, R.P., Limbach, P.A., and Woodson, S.A. (2013). In vivo X-ray footprinting of pre-30S ribosomes reveals chaperone-dependent remodeling of late assembly intermediates. Mol Cell 52, 506–516.

Daigle, D.M., Rossi, L., Berghuis, A.M., Aravind, L., Koonin, E.V., and Brown, E.D. (2002). YjeQ, an essential, conserved, uncharacterized protein from Escherichia coli, is an unusual GTPase with circularly permuted G-motifs and marked burst kinetics. Biochemistry 41, 11109–11117.

Datsenko, K.A., and Wanner, B.L. (2000). One-step inactivation of chromosomal genes in Escherichia coli K-12 using PCR products. Proc Natl Acad Sci U S A 97, 6640–6645.

Datta, P.P., Wilson, D.N., Kawazoe, M., Swami, N.K., Kaminishi, T., Sharma, M.R., Booth, T.M., Takemoto, C., Fucini, P., Yokoyama, S., et al. (2007). Structural aspects of RbfA action during small ribosomal subunit assembly. Mol Cell 28, 434–445.

Datta, S., Costantino, N., and Court, D.L. (2006). A set of recombineering plasmids for gram-negative bacteria. Gene 379, 109–115.

Davis, J.H., Tan, Y.Z., Carragher, B., Potter, C.S., Lyumkis, D., and Williamson, J.R. (2016). Modular Assembly of the Bacterial Large Ribosomal Subunit. Cell 167, 1610–1622 e1615.

Deutsch, E.W., Shteynberg, D., Lam, H., Sun, Z., Eng, J.K., Carapito, C., von Haller, P.D., Tasman, N., Mendoza, L., Farrah, T., et al. (2010). Trans-Proteomic Pipeline supports and improves analysis of electron transfer dissociation data sets. Proteomics 10, 1190–1195.

Emsley, P., and Cowtan, K. (2004). Coot: model-building tools for molecular graphics. Acta Crystallogr D Biol Crystallogr 60, 2126–2132.

Emsley, P., Lohkamp, B., Scott, W.G., and Cowtan, K. (2010). Features and development of Coot. Acta Crystallogr D Biol Crystallogr 66, 486–501.

Eng, J.K., Jahan, T.A., and Hoopmann, M.R. (2013). Comet: an open-source MS/MS sequence database search tool. Proteomics 13, 22–24.

Escher, C., Reiter, L., MacLean, B., Ossola, R., Herzog, F., Chilton, J., MacCoss, M.J., and Rinner, O. (2012). Using iRT, a normalized retention time for more targeted measurement of peptides. Proteomics 12, 1111–1121.

Ghosal, A., Babu, V.M.P., and Walker, G.C. (2018). Elevated Levels of Era GTPase Improve Growth, 16S rRNA Processing, and 70S Ribosome Assembly of Escherichia coli Lacking Highly Conserved Multifunctional YbeY Endoribonuclease. J Bacteriol 200.

Guo, Q., Goto, S., Chen, Y., Feng, B., Xu, Y., Muto, A., Himeno, H., Deng, H., Lei, J., and Gao, N. (2013). Dissecting the in vivo assembly of the 30S ribosomal subunit reveals the role of RimM and general features of the assembly process. Nucleic Acids Res 41, 2609–2620.

Guo, Q., Yuan, Y., Xu, Y., Feng, B., Liu, L., Chen, K., Sun, M., Yang, Z., Lei, J., and Gao, N. (2011). Structural basis for the function of a small GTPase RsgA on the 30S ribosomal subunit maturation revealed by cryoelectron microscopy. Proc Natl Acad Sci U S A 108, 13100–13105.

Hang, J.Q., and Zhao, G. (2003). Characterization of the 16S rRNA- and membrane-binding domains of Streptococcus pneumoniae Era GTPase: structural and functional implications. Eur J Biochem 270, 4164–4172.

Hao, Y., Bohon, J., Hulscher, R., Rappe, M.C., Gupta, S., Adilakshmi, T., and Woodson, S.A. (2018). Time-Resolved Hydroxyl Radical Footprinting of RNA with X-Rays. Curr Protoc Nucleic Acid Chem 73, e52.

Hao, Y., and Kieft, J.S. (2014). Diverse self-association properties within a family of phage packaging RNAs. RNA 20, 1759–1774.

Himeno, H., Hanawa-Suetsugu, K., Kimura, T., Takagi, K., Sugiyama, W., Shirata, S., Mikami, T., Odagiri, F., Osanai, Y., Watanabe, D., et al. (2004). A novel GTPase activated by the small subunit of ribosome. Nucleic Acids Res 32, 5303–5309.

Inoue, K., Alsina, J., Chen, J., and Inouye, M. (2003). Suppression of defective ribosome assembly in a rbfA deletion mutant by overexpression of Era, an essential GTPase in Escherichia coli. Mol Microbiol 48, 1005–1016.

Ji, X. (2016). Structural insights into cell cycle control by essential GTPase Era. Postepy Biochem 62, 335–342.

Johnstone, B.H., Handler, A.A., Chao, D.K., Nguyen, V., Smith, M., Ryu, S.Y., Simons, E.L., Anderson, P.E., and Simons, R.W. (1999). The widely conserved Era G-protein contains an RNA-binding domain required for Era function in vivo. Mol Microbiol 33, 1118–1131.

Jomaa, A., Jain, N., Davis, J.H., Williamson, J.R., Britton, R.A., and Ortega, J. (2014). Functional domains of the 50S subunit mature late in the assembly process. Nucleic Acids Res 42, 3419–3435.

Jomaa, A., Stewart, G., Martin-Benito, J., Zielke, R., Campbell, T.L., Maddock, J.R., Brown, E.D., and Ortega, J. (2011a). Understanding ribosome assembly: the structure of in vivo assembled immature 30S subunits revealed by cryo-electron microscopy. RNA 17, 697709.

Jomaa, A., Stewart, G., Mears, J.A., Kireeva, I., Brown, E.D., and Ortega, J. (2011b). Cryo-electron microscopy structure of the 30S subunit in complex with the YjeQ biogenesis factor. RNA 17, 2026–2038.

Karbstein, K. (2013). Quality control mechanisms during ribosome maturation. Trends Cell Biol 23, 242–250.

Kimanius, D., Forsberg, B.O., Scheres, S.H., and Lindahl, E. (2016). Accelerated cryo-EM structure determination with parallelisation using GPUs in RELION-2. eLife 5.

Lam, H., Deutsch, E.W., Eddes, J.S., Eng, J.K., King, N., Stein, S.E., and Aebersold, R. (2007). Development and validation of a spectral library searching method for peptide identification from MS/MS. Proteomics 7, 655–667.

Lauber, M.A., Rappsilber, J., and Reilly, J.P. (2012). Dynamics of ribosomal protein S1 on a bacterial ribosome with cross-linking and mass spectrometry. Mol Cell Proteomics 11, 1965–1976.

Leipe, D.D., Wolf, Y.I., Koonin, E.V., and Aravind, L. (2002). Classification and evolution of P-loop GTPases and related ATPases. J Mol Biol 317, 41–72.

Leong, V., Kent, M., Jomaa, A., and Ortega, J. (2013). Escherichia coli rimM and yjeQ null strains accumulate immature 30S subunits of similar structure and protein complement. RNA 19, 789–802.

Link, A.J., Phillips, D., and Church, G.M. (1997). Methods for generating precise deletions and insertions in the genome of wild-type Escherichia coli: application to open reading frame characterization. J Bacteriol 179, 6228–6237.

Lopez-Alonso, J.P., Kaminishi, T., Kikuchi, T., Hirata, Y., Iturrioz, I., Dhimole, N., Schedlbauer, A., Hase, Y., Goto, S., Kurita, D., et al. (2017). RsgA couples the maturation state of the 30S ribosomal decoding center to activation of its GTPase pocket. Nucleic Acids Res 45, 6945–6959.

Lu, Q., and Inouye, M. (1998). The gene for 16S rRNA methyltransferase (ksgA) functions as a multicopy suppressor for a cold-sensitive mutant of era, an essential RAS-like GTP-binding protein in Escherichia coli. J Bacteriol 180, 5243–5246.

MacLean, B., Tomazela, D.M., Shulman, N., Chambers, M., Finney, G.L., Frewen, B., Kern, R., Tabb, D.L., Liebler, D.C., and MacCoss, M.J. (2010). Skyline: an open source document editor for creating and analyzing targeted proteomics experiments. Bioinformatics 26, 966–968.

March, P.E., Lerner, C.G., Ahnn, J., Cui, X., and Inouye, M. (1988). The Escherichia coli Ras-like protein (Era) has GTPase activity and is essential for cell growth. Oncogene 2, 539–544.

Minkovsky, N., Zarimani, A., Chary, V.K., Johnstone, B.H., Powell, B.S., Torrance, P.D., Court, D.L., Simons, R.W., and Piggot, P.J. (2002). Bex, the Bacillus subtilis homolog of the essential Escherichia coli GTPase Era, is required for normal cell division and spore formation. J Bacteriol 184, 6389–6394.

Mulder, A.M., Yoshioka, C., Beck, A.H., Bunner, A.E., Milligan, R.A., Potter, C.S., Carragher, B., and Williamson, J.R. (2010). Visualizing ribosome biogenesis: parallel assembly pathways for the 30S subunit. Science 330, 673–677.

Ni, X., Davis, J.H., Jain, N., Razi, A., Benlekbir, S., McArthur, A.G., Rubinstein, J.L., Britton, R.A., Williamson, J.R., and Ortega, J. (2016). YphC and YsxC GTPases assist the maturation of the central protuberance, GTPase associated region and functional core of the 50S ribosomal subunit. Nucleic Acids Res 44, 8442–8455.

Pettersen, E.F., Goddard, T.D., Huang, C.C., Couch, G.S., Greenblatt, D.M., Meng, E.C., and Ferrin, T.E. (2004). UCSF Chimera--a visualization system for exploratory research and analysis. J Comput Chem 25, 1605–1612.

Powers, T., and Noller, H.F. (1990). Dominant lethal mutations in a conserved loop in 16S rRNA. Proc Natl Acad Sci U S A 87, 1042–1046.

Prossliner, T., Skovbo Winther, K., Sorensen, M.A., and Gerdes, K. (2018). Ribosome Hibernation. Annu Rev Genet 52, 321–348.

Razi, A., Guarne, A., and Ortega, J. (2017). The cryo-EM structure of YjeQ bound to the 30S subunit suggests a fidelity checkpoint function for this protein in ribosome assembly. Proc Natl Acad Sci U S A 114, E3396–E3403.

Rosenthal, P.B., and Henderson, R. (2003). Optimal determination of particle orientation, absolute hand, and contrast loss in single-particle electron cryomicroscopy. J Mol Biol 333, 721–745.

Sashital, D.G., Greeman, C.A., Lyumkis, D., Potter, C.S., Carragher, B., and Williamson, J.R. (2014). A combined quantitative mass spectrometry and electron microscopy analysis of ribosomal 30S subunit assembly in E. coli. eLife 3.

Sato, T., Wu, J., and Kuramitsu, H. (1998). The sgp gene modulates stress responses of Streptococcus mutans: utilization of an antisense RNA strategy to investigate essential gene functions. FEMS Microbiol Lett 159, 241–245.

Sayed, A., Matsuyama, S., and Inouye, M. (1999). Era, an essential Escherichia coli small G-protein, binds to the 30S ribosomal subunit. Biochem Biophys Res Commun 264, 51–54.

Scheres, S.H. (2012). RELION: implementation of a Bayesian approach to cryo-EM structure determination. J Struct Biol 180, 519–530.

Schuwirth, B.S., Borovinskaya, M.A., Hau, C.W., Zhang, W., Vila-Sanjurjo, A., Holton, J.M., and Cate, J.H. (2005). Structures of the bacterial ribosome at 3.5 A resolution. Science 310, 827–834.

Shajani, Z., Sykes, M.T., and Williamson, J.R. (2011). Assembly of bacterial ribosomes. Annu Rev Biochem 80, 501–526.

Sharma, M.R., Barat, C., Wilson, D.N., Booth, T.M., Kawazoe, M., Hori-Takemoto, C., Shirouzu, M., Yokoyama, S., Fucini, P., and Agrawal, R.K. (2005). Interaction of Era with the 30S ribosomal subunit implications for 30S subunit assembly. Mol Cell 18, 319–329.

Shteynberg, D., Deutsch, E.W., Lam, H., Eng, J.K., Sun, Z., Tasman, N., Mendoza, L., Moritz, R.L., Aebersold, R., and Nesvizhskii, A.I. (2011). iProphet: multi-level integrative analysis of shotgun proteomic data improves peptide and protein identification rates and error estimates. Mol Cell Proteomics 10, M111 007690.

Strunk, B.S., Loucks, C.R., Su, M., Vashisth, H., Cheng, S., Schilling, J., Brooks, C.L., 3rd, Karbstein, K., and Skiniotis, G. (2011). Ribosome assembly factors prevent premature translation initiation by 40S assembly intermediates. Science 333, 1449–1453.

Strunk, B.S., Novak, M.N., Young, C.L., and Karbstein, K. (2012). A translation-like cycle is a quality control checkpoint for maturing 40S ribosome subunits. Cell 150, 111–121.

Sturn, A., Quackenbush, J., and Trajanoski, Z. (2002). Genesis: cluster analysis of microarray data. Bioinformatics 18, 207–208.

Sykes, M.T., and Williamson, J.R. (2009). A Complex Assembly Landscape for the 30S Ribosomal Subunit. Annu Rev Biophys 38, 197–215.

Takiff, H.E., Chen, S.M., and Court, D.L. (1989). Genetic analysis of the rnc operon of Escherichia coli. J Bacteriol 171, 2581–2590.

Talkington, M.W., Siuzdak, G., and Williamson, J.R. (2005). An assembly landscape for the 30S ribosomal subunit. Nature 438, 628–632.

Tang, G., Peng, L., Baldwin, P.R., Mann, D.S., Jiang, W., Rees, I., and Ludtke, S.J. (2007). EMAN2: an extensible image processing suite for electron microscopy. J Struct Biol 157, 38–46.

Terwilliger, T.C., Sobolev, O.V., Afonine, P.V., and Adams, P.D. (2018). Automated map sharpening by maximization of detail and connectivity. Acta Crystallogr D Struct Biol 74, 545–559.

Thurlow, B., Davis, J.H., Leong, V., T, F.M., Williamson, J.R., and Ortega, J. (2016). Binding properties of YjeQ (RsgA), RbfA, RimM and Era to assembly intermediates of the 30S subunit. Nucleic Acids Res 44, 9918–9932.

Traub, P., and Nomura, M. (1968). Structure and function of Escherichia coli ribosomes. I. Partial fractionation of the functionally active ribosomal proteins and reconstitution of artificial subribosomal particles. J Mol Biol 34, 575–593.

Traub, P., and Nomura, M. (1969a). Structure and function of Escherichia coli ribosomes. VI. Mechanism of assembly of 30 s ribosomes studied in vitro. J Mol Biol 40, 391–413.

Traub, P., and Nomura, M. (1969b). Studies on the assembly of ribosomes in vitro. Cold Spring Harb Symp Quant Biol 34, 63–67.

Tu, C., Zhou, X., Tropea, J.E., Austin, B.P., Waugh, D.S., Court, D.L., and Ji, X. (2009). Structure of ERA in complex with the 3’ end of 16S rRNA: implications for ribosome biogenesis. Proc Natl Acad Sci U S A 106, 14843–14848.

Vargas, J., Alvarez-Cabrera, A.L., Marabini, R., Carazo, J.M., and Sorzano, C.O. (2014). Efficient initial volume determination from electron microscopy images of single particles. Bioinformatics 30, 2891–2898.

Yang, Z., Guo, Q., Goto, S., Chen, Y., Li, N., Yan, K., Zhang, Y., Muto, A., Deng, H., Himeno, H., et al. (2014). Structural insights into the assembly of the 30S ribosomal subunit in vivo: functional role of S5 and location of the 17S rRNA precursor sequence. Protein & cell 5, 394–407.

Zhang, K. (2016). Gctf: Real-time CTF determination and correction. J Struct Biol 193, 1–12.

Zheng, S.Q., Palovcak, E., Armache, J.P., Verba, K.A., Cheng, Y., and Agard, D.A. (2017). MotionCor2: anisotropic correction of beam-induced motion for improved cryo-electron microscopy. Nat Methods 14, 331–332.

Zundel, M.A., Basturea, G.N., and Deutscher, M.P. (2009). Initiation of ribosome degradation during starvation in Escherichia coli. RNA 15, 977–983.

