## Supplemental Information for "Role of Era in Assembly and Homeostasis of the Ribosomal Small Subunit"

[1]Department of Anatomy and Cell Biology, McGill University, Montreal, Quebec, Canada. [2]Department of Molecular Biology, [3]Department of Chemistry and The Skaggs Institute for Chemical Biology, The Scripps Research Institute, La Jolla, CA, USA. [4]Department of Biology, Massachusetts Institute of Technology, Cambridge, MA. [5]T.C. Jenkins Department of Biophysics, Johns Hopkins University, Baltimore, MD, USA. [6]Department of Biochemistry and Biomedical Sciences, McMaster University, Hamilton, Ontario, Canada. [7]Department of Molecular Virology and Microbiology, [8]Center for Metagenomics and Microbiome Research, Baylor College of Medicine, Houston, TX, USA. [9]Department of Biochemistry, McGill University, Montreal, Quebec, Canada.

**This supplement contains:**

Supplemental Results
Supplemental Figures S1 to S8
Supplemental Table 1
Supplemental References

### SUPPLEMENTAL RESULTS

#### Era depleted cells exhibit slow growth and accumulate unprocessed rRNA

To generate the Era-depleted strain, an *era* gene was first inserted into the bacterial chromosome under the control of the *ara*BAD arabinose inducible promoter. Subsequently, the native *era* gene was removed. Replacement of the *araBAD* genes with *era* and the precise deletion of the native *era* gene at its native locus was confirmed by PCR screening (Figure S1A and S1B and Star Methods). Consistent with previous literature (Inoue et al., 2003; Lerner and Inouye, 1991), the growth of the Era-depleted strain in the absence of arabinose was drastically reduced relative to the wild type strains. When the Era-depleted strain was grown in the presence of arabinose the culture reached the stationary phase at a lower optical density compared to the wild type strain (Figure S1B, left panel).  The slower growth of this strain in the presence of arabinose was also apparent in solid media (Figure S1C, right panel) and removing the inducer led to a complete disappearance of colonies in all dilutions of the assay.

To ensure that other phenotypical characteristics of the newly generated Era-depleted strain were similar to previously characterized strains with a similar genetic modification, we also measured the level of unprocessed 16S rRNA under Era depletion conditions by quantitative PCR (qPCR). Compared to parental cells, we found that this strain under Era depletion conditions had 9.6±0.51 folds higher 17S/16S rRNA molecules containing the 115 nucleotides that comprise the 5' precursor sequence and 2.7±0.57 folds higher fragments with the 33 extra bases in the 3' precursor sequence. These results are in agreement with observations in other *E. coli* Era-depleted strains (Inoue et al., 2003).

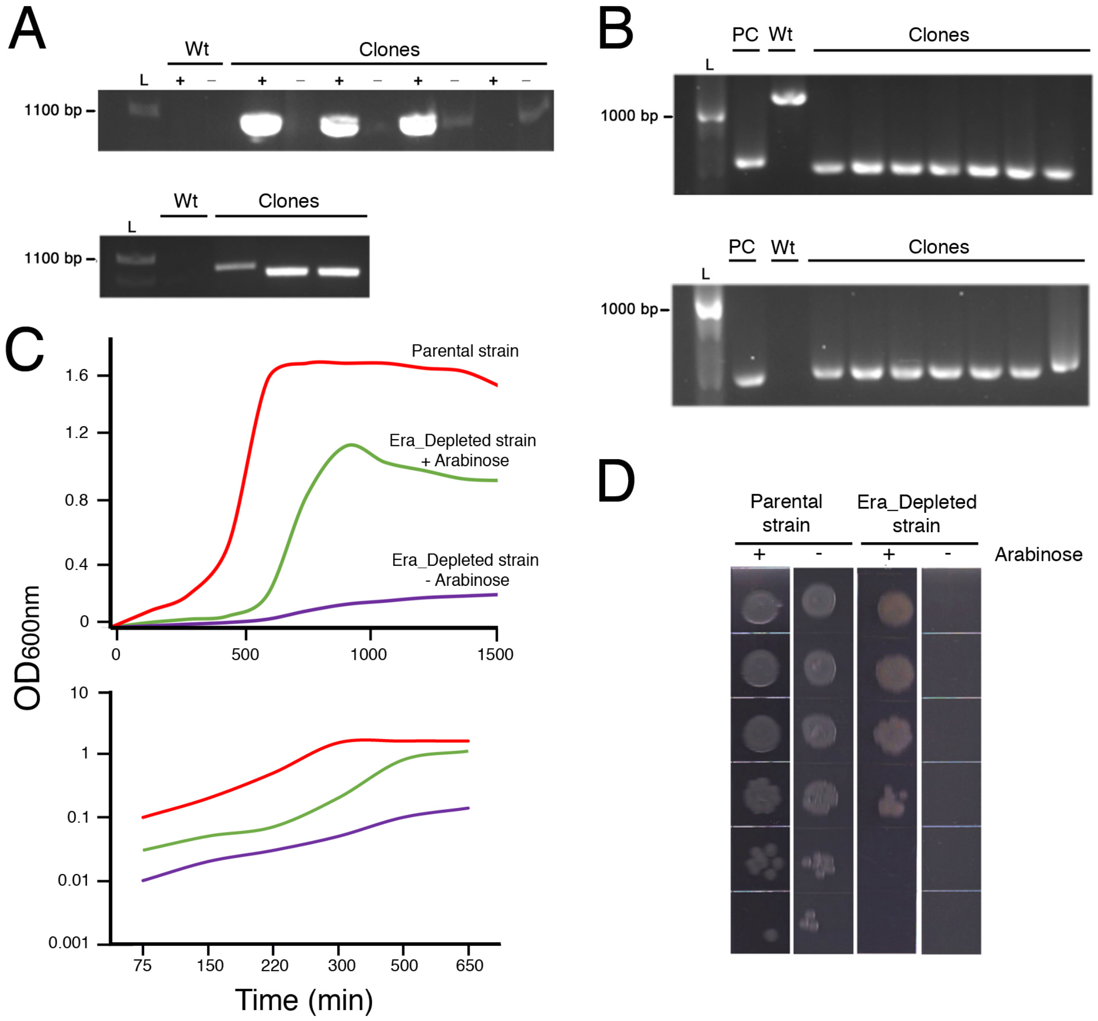


**Figure S1. Generation and phenotype of the Era-depleted strain.** (A) PCR screening results using Ara_up-R and Era_down-F oligonucleotides (+) (insertion in correct orientation) or Ara_up-R and Era_up-R oligonucleotides (-) (insertion in wrong orientation) (top panel) or kan_int and ara_int oligonucletodies (confirms insertion) (bottom panel). These PCR results demonstrate insertion of the double digested (*NotI* and *PsiI*) linearized DNA fragment from the pBS-*araBAD*flank*erakan* into the chromosome of *E.coli* BW25113 cells to generate an *araBAD*::*era* strain. (B) PCR screening results using Apra_int-F and EraKO_confirm-R primers (top panel) and apra_int-R and EraKO_confirm-F (bottom panel) to confirm the precise knockout of *era* and insertion of *apramycin*$^{r}$ at the *era* locus in *araBAD*::*era E.coli* BW25113 cells to generate *araBAD*::*era*, *era*::*apr*$^r$. The top panel show a PCR amplicon for the wild type (Wt) strain using the EraKO_confirm-F and EraKO_confirm-R oligonucleotides with an expected product of ~1100 bp. The bottom panel shows the PCR results for the wild type (Wt) using the apra_int-R and EraKO_confirm-F oligonucleotides, which is not expected to produce an amplicon of the native *era* locus. A positive control (PC) was used to ensure optimal PCR amplification for the apramycin$^r$ cassette. (C) Growth profiles of the parental and Era depleted strain in LB liquid media (top panel). The growth profile of the Era depleted strain is shown for the cells grown in the liquid media with and without 1% arabinose. The exponential growth phase of the three curves was plotted in a semi-log plot for easy comparison of the growth rates. (D) Dilution plating experiments of saturated cultures of parental and Era depleted strains in LB agar plates with and without 1% arabinose. Spots represent 10-fold dilution increment of the saturated culture. Figure S1 related to Figure 1.

A

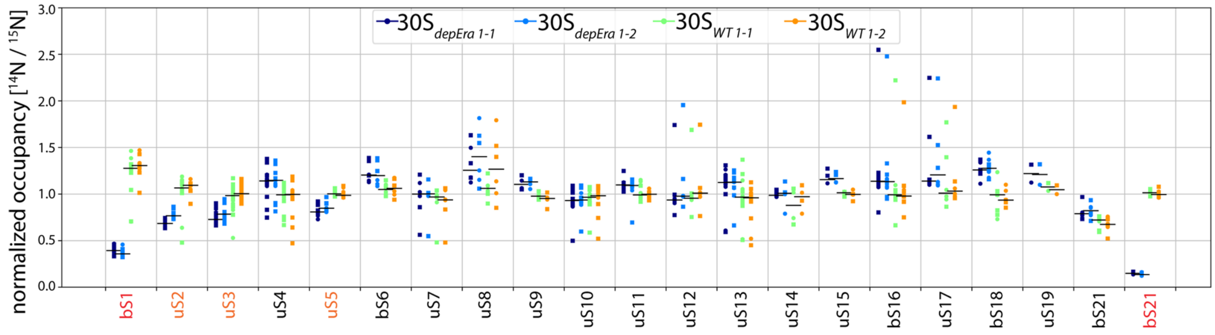


B

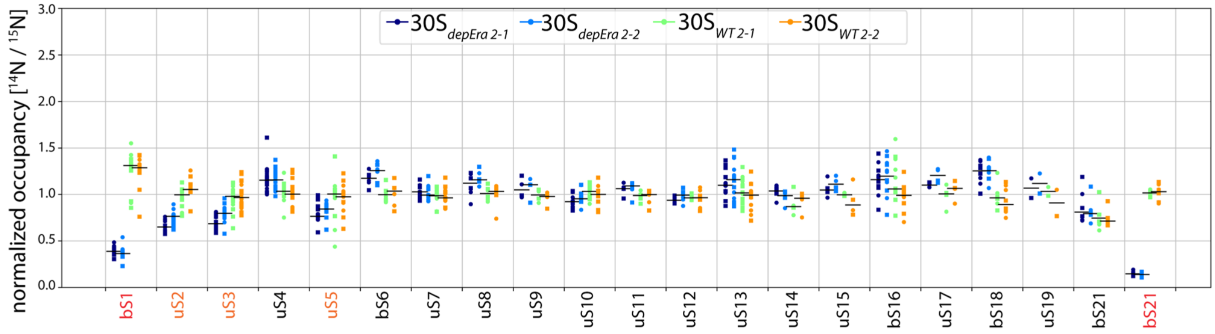


**Figure S2. Protein composition of the $30S_{Era\text{-}depleted}$ particles.** (A) Dot plots showing occupancy of the r-proteins in the 30S particles purified from the parental strain ($30S_{WT}$) and the Era depleted strain grown under (-arabinose) conditions ($30S_{depEra}$). Occupancy is calculated as the $^{14}N/^{15}N$ ratio /median ($^{14}N/^{15}N$ for all small subunit proteins), where $^{14}N$ is for the experimental 30S and $^{15}N$ is for a purified 70S particle from wild type cells. Data points shown as circles are from quantifying the MS1 spectra, and data points shown as squares are from quantifying the MS/MS product ion spectra. Proteins in group 1 (stoichiometric), group 2 (intermediate occupancy) and group 3 (low occupancy) are labeled in black, orange and red, respectively. (B) Experimental replica of the experiment shown in (A). Figure S2 related to Figure 1.

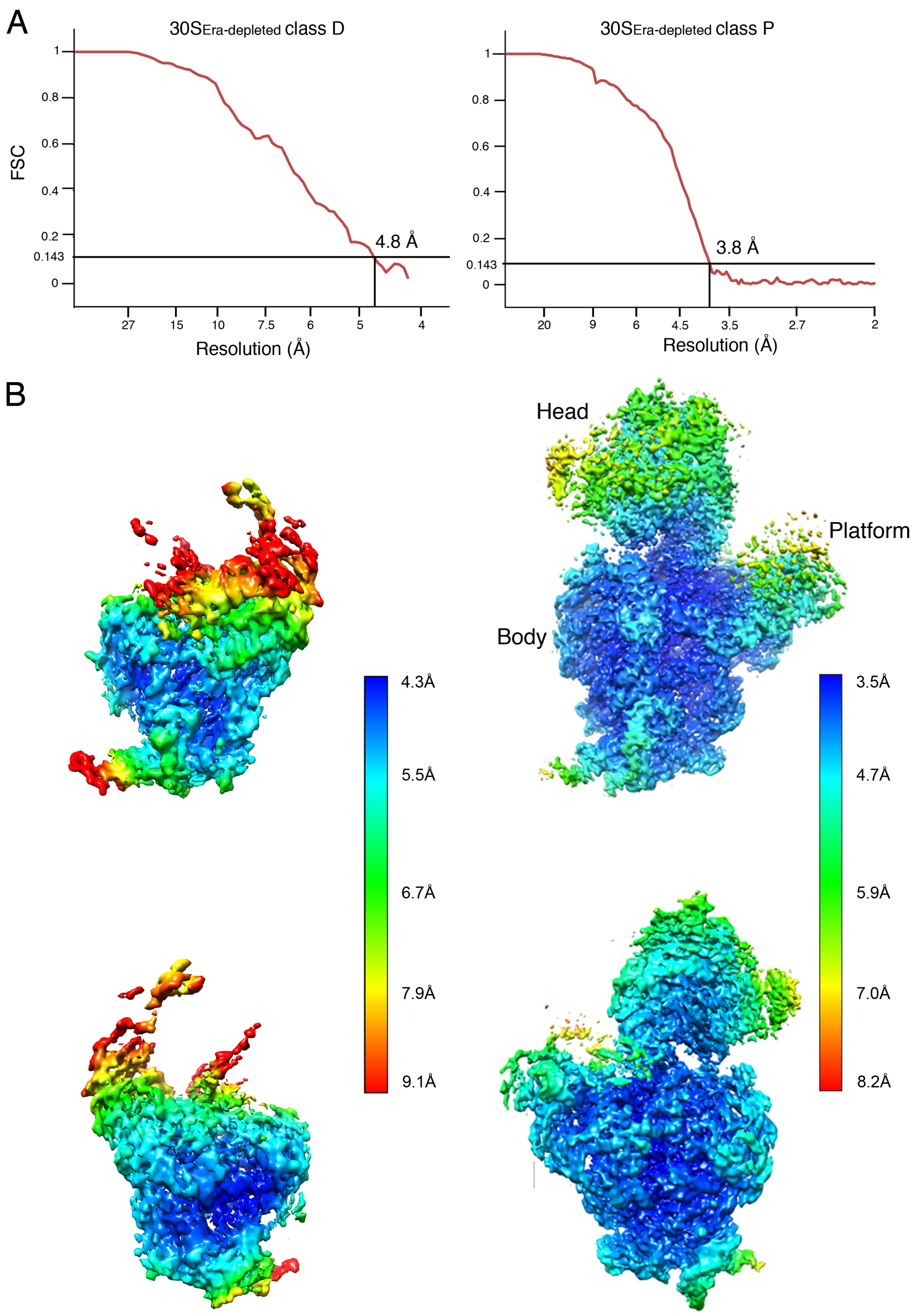


**Figure S3. Resolution analysis of the cryo-EM structures obtained for the $30S_{Era\text{-}depleted}$ particles.** (A) Fourier shell correlation (FSC) plots for the cryo-EM maps of the $30S_{Era\text{-}depleted}$ classes D and P. Resolution is reported using a FSC threshold of 0.143. (B) Local resolution analysis of the cryo-EM maps for classes D and P. Figure S3 related to Figure 3.

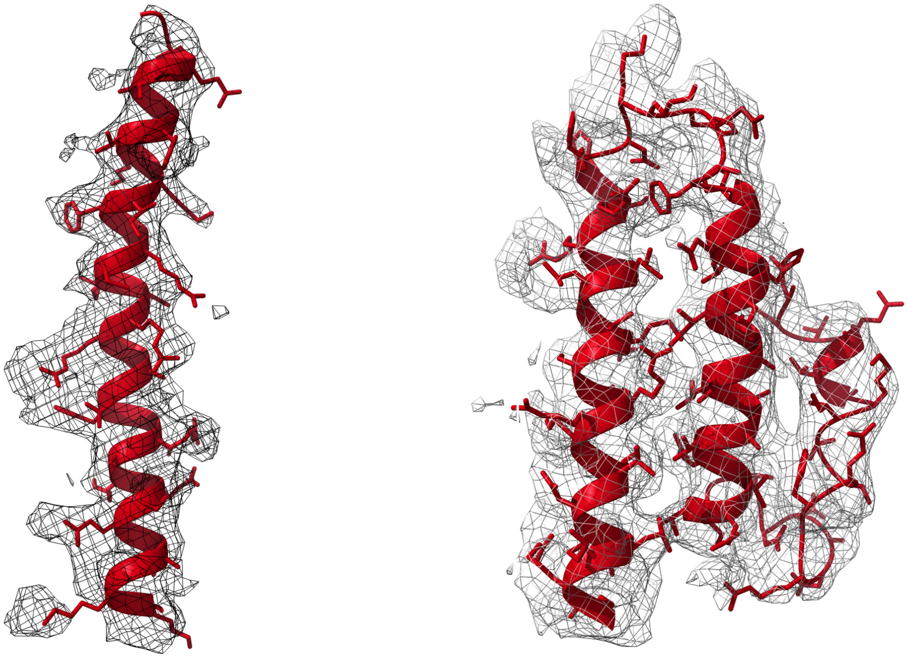


**Figure S4. Fitting of the atomic model of the $30S_{Era\text{-}depleted}$ structure (class P) into the cryo-EM map.** Illustrative densities for r-proteins bS20 and uS15 in the cryo-EM map and the corresponding atomic model of the $30S_{Era\text{-}depleted}$ (class P) built for these regions. Figure S4 related to Figure 3.

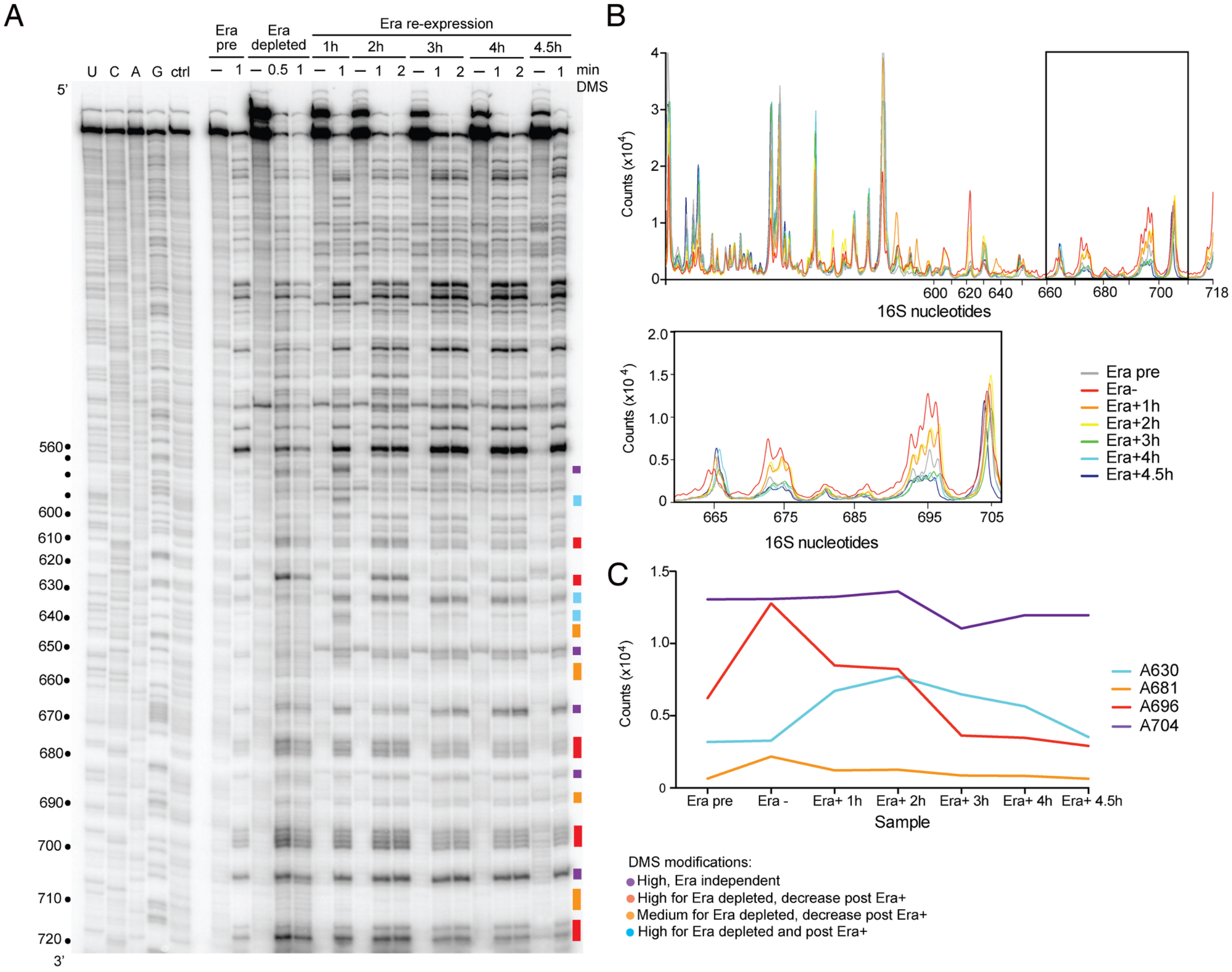


**Figure S5. Quantification of *in vivo* DMS footprinting after Era depletion.** (A) Sequencing gel similar to Figure 4B except that the gel was run for 4 h. DMS modified nucleotides are highlighted on the right and colored based on their modification patterns. (B) Band intensities of the primer extension gel shown in panel A as indicated in the key. The region corresponding to nucleotides 660 to 706 (h23) is highlighted with a dark gray bar and expanded in the lower panel. (C) Relative DMS modification of representative nucleotides with different Era modification patterns. The DMS modification patterns of different nucleotides are graphed with the same color scheme as panel A. Figure S5 related to Figure 4.

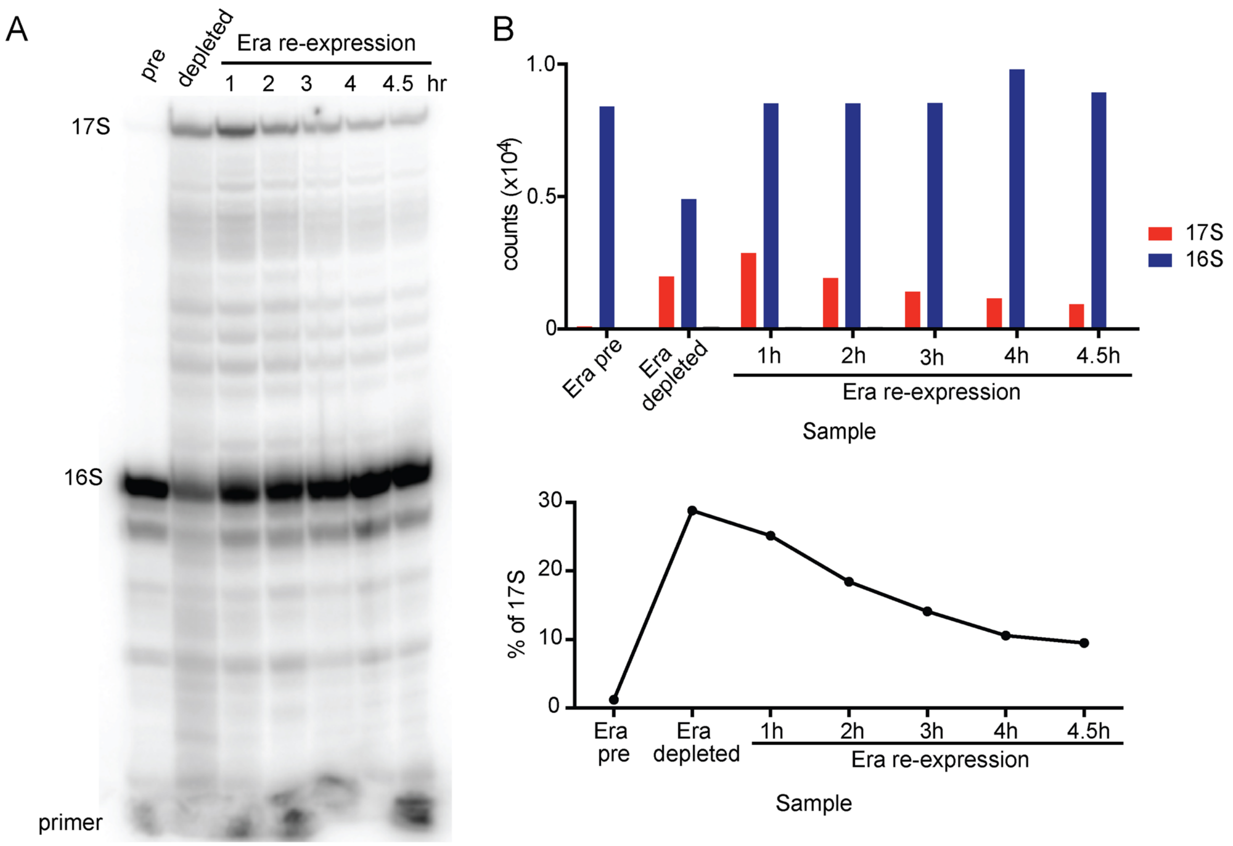


**Figure S6. Era depletion impairs pre-rRNA processing.** (A) Primer extension of DMS-modified RNA after Era depletion and restoration with primer 46. Products corresponding to the mature 16S 5′ end and the 17S precursor are indicated. Samples as in Figure 4. (B) Top; intensities of 17S (red) and 16S bands (blue). Bottom; percentage of 17S relative to total rRNA. Figure S6 related to Figure 4.

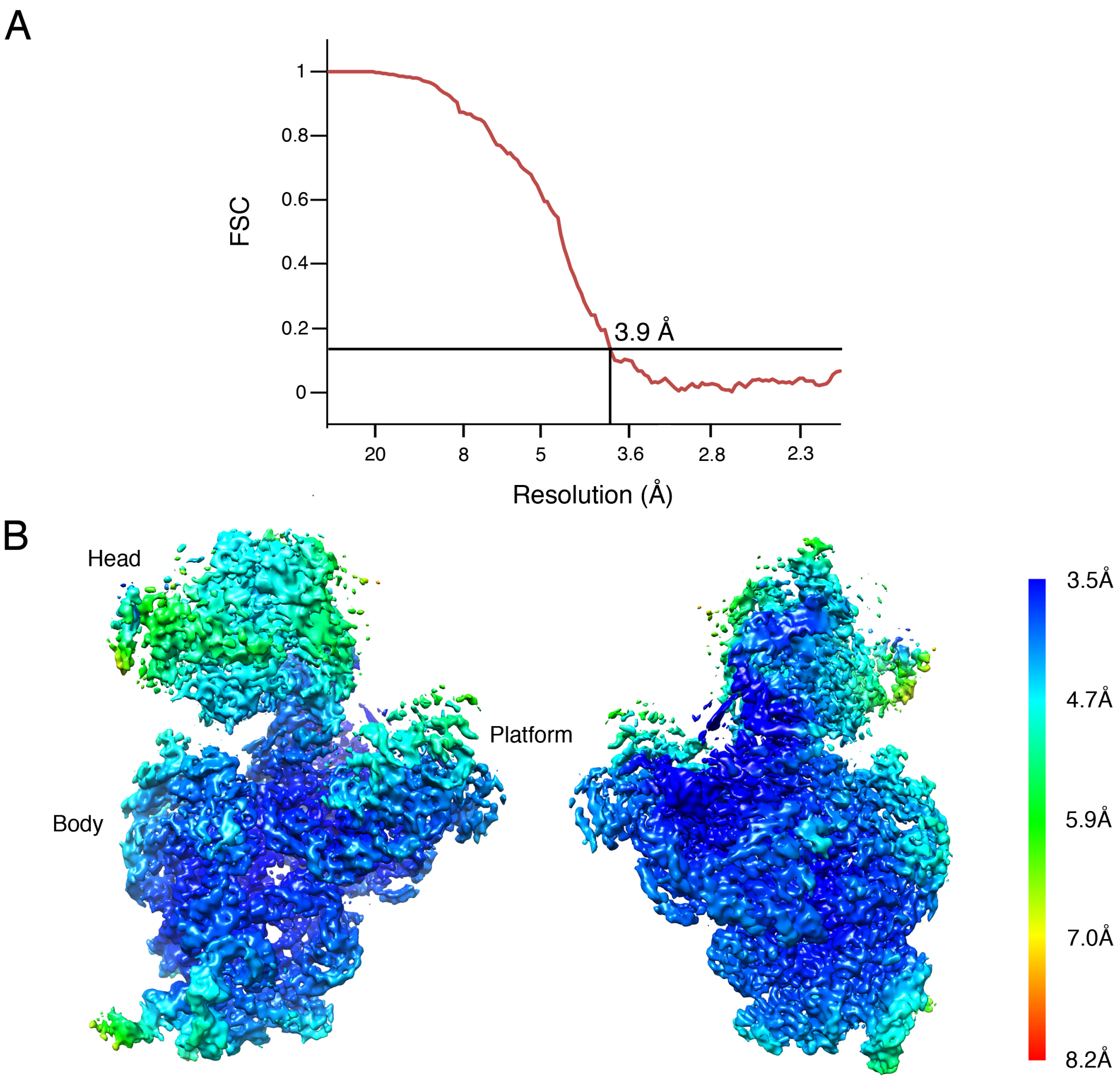


**Figure S7. Resolution analysis of the 30S+Era complex.** (A) Average resolution for the cryo-EM maps of the Era-treated 30S particles was estimated by gold-standard Fourier shell correlation. Resolution estimation is reported using a FSC threshold value of 0.143. (B) Local resolution analysis for the cryo-EM map of the 30S+Era complex. Figure S7 related to Figure 5.

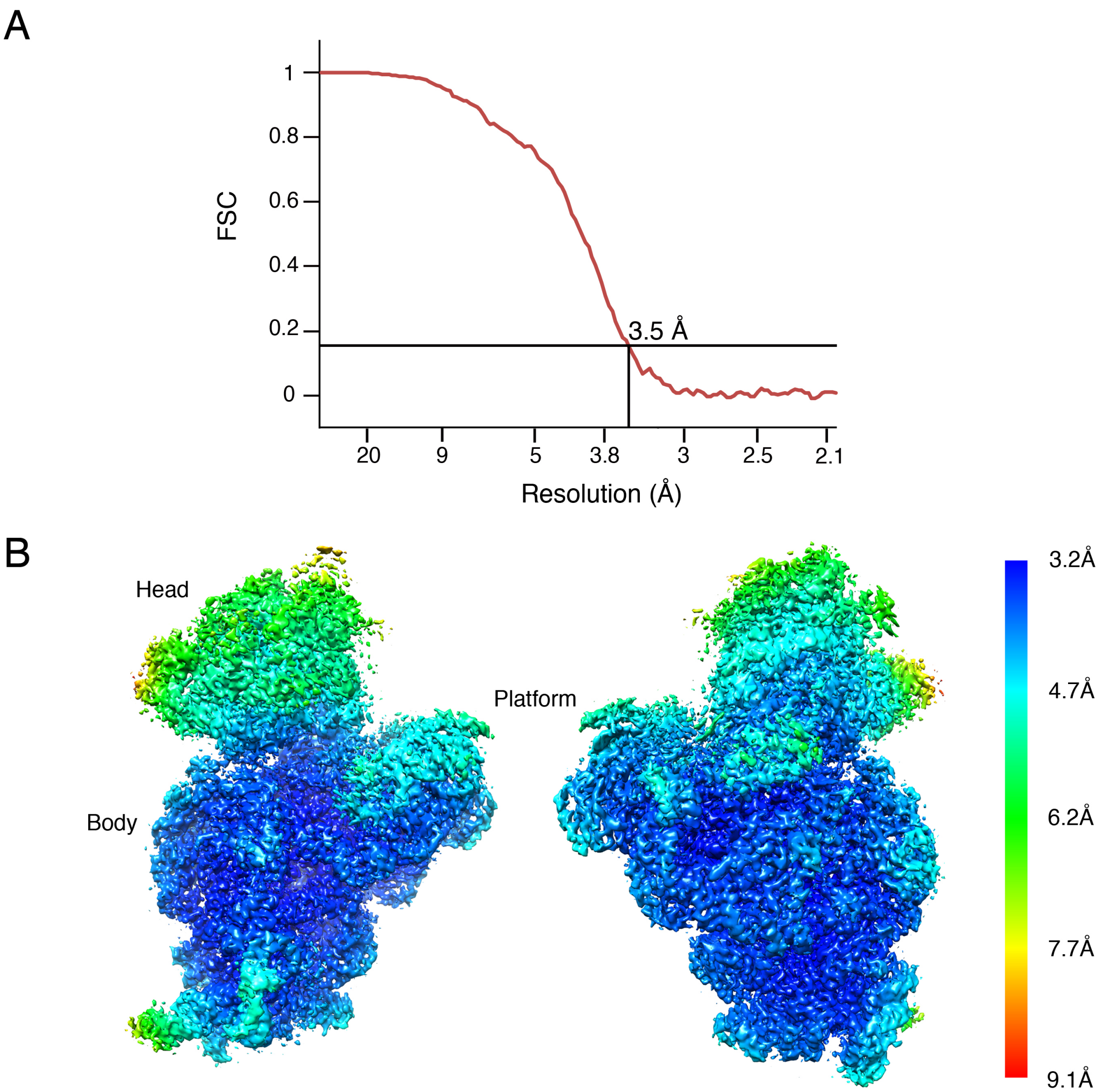


**Figure S8. Resolution analysis of the 30S+Era+YjeQ complex.** (A) Fourier shell correlation plot for the cryo-EM map of the Era+YjeQ-treated 30S particles. Resolution estimation is reported using a FSC threshold value of 0.143. (B) Local resolution analysis performed for the cryo-EM map of the 30S+Era+YjeQ complex. Figure S8 related to Figure 6.

### SUPPLEMENTAL TABLES

**Supplemental Table 1. Cryo-EM data collection, refinement and structure validation.**

| | $30S_{Era\text{-}depleted}$ | Era-treated 30S | Era+YjeQ treated 30S |
|---|---|---|---|
| **Data collection** | | | |
| Microscope | Titan Krios | Titan Krios | Titan Krios |
| Detector | Falcon II | Falcon II | Falcon II |
| Magnification | 75,000x | 75,000x | 75,000x |
| Voltage (kV) | 300 | 300 | 300 |
| Total electron dose ($e^-$/Å | 35 | 46 | 28 |
| Defocus range (µm) | -1.25 to -2.75 | -1.25 to -2.75 | -1.25 to -2.75 |
| Pixel size (Å/px) | 1.073 | 1.073 | 1.073 |
| **Reconstruction and refinement** | | | |
| Particles | 233,149 | 250,896 | 195,841 |
| Map sharpening B factor | -176 | -152 | -125 |
| Resolution (Å) | 3.8 | 3.9 | 3.5 |
| FSC Threshold | 0.143 | 0.143 | 0.143 |
| **Model composition** | | | |
| RNA chains | 1 | NA | NA |
| Protein chains | 16 | NA | NA |
| **Model validation** | | | |
| Ramachandran outliers | 0.06% | NA | NA |
| Ramachandran favored | 86.31% | NA | NA |
| MolProbity score | 2.07 | NA | NA |
| Clashscore | 7.99 | NA | NA |
| Rotamer Outliers | 0.41% | NA | NA |

### SUPPLEMENTARY REFERENCES

Inoue, K., Alsina, J., Chen, J., and Inouye, M. (2003). Suppression of defective ribosome assembly in a rbfA deletion mutant by overexpression of Era, an essential GTPase in Escherichia coli. Mol Microbiol *48*, 1005-1016.

Lerner, C.G., and Inouye, M. (1991). Pleiotropic changes resulting from depletion of Era, an essential GTP-binding protein in Escherichia coli. Mol Microbiol *5*, 951-957.
